# Nuclear Pif1 is Post Translationally Modified and Regulated by Lysine Acetylation

**DOI:** 10.1101/2020.07.06.189761

**Authors:** Onyekachi E. Ononye, Christopher W. Sausen, Lata Balakrishnan, Matthew L. Bochman

**Affiliations:** Department of Biology, School of Science, Indiana University Purdue University Indianapolis, Indianapolis, Indiana, USA; Molecular and Cellular Biochemistry Department, Indiana University, Bloomington, Indiana, USA

**Keywords:** Pif1 Helicase, Lysine Acetylation, Piccolo NuA4, Rpd3, DNA Replication and Repair

## Abstract

In *S. cerevisiae*, the Pif1 helicase functions to impact both nuclear and mitochondrial DNA replication and repair processes. Pif1 is a 5’-3’ helicase, which preferentially unwinds RNA-DNA hybrids and resolves G-quadruplex structures. Further, regulation of Pif1 by phosphorylation negatively impacts its interaction with telomerase during double strand break repair. Here, we report that in addition to phosphorylation, Pif1 is also modified by lysine acetylation, which influences both its cellular and core biochemical activities. Using Pif1 overexpression toxicity assays, we determined that the acetyltransferase NuA4 (Esa1) and deacetylase Rpd3 are primarily responsible for dynamically acetylating nuclear Pif1. Mass spectrometry analysis revealed that Pif1 was modified throughout the protein’s sequence on the N-terminus (K118, K129), helicase domain (K525, K639, K725), and C-terminus (K800). Acetylation of Pif1 exacerbated its overexpression toxicity phenotype, which was alleviated upon deletion of its N-terminus. Biochemical assays demonstrated that acetylation of Pif1 stimulated its helicase activity, while maintaining its substrate preferences. Additionally, both the ATPase and DNA binding activities of Pif1 were stimulated upon acetylation. Limited proteolysis assays indicate that acetylation of Pif1 induces a conformational change that may account for its altered enzymatic properties. We propose an acetylation-based model for the regulation of Pif1 activities, addressing how this post translational modification can influence its role as a key player in a multitude of DNA transactions vital to the maintenance of genome integrity.

## INTRODUCTION

At the core of all cellular transactions, such as replication, repair, and transcription, is the need for biological machines to gain access to the genetic information stored within the DNA duplex [1]. Along with chromatin remodeling, access to the DNA is provided through either the active or passive action of helicases, which function to unwind double-stranded DNA (dsDNA) into its complementary single strands. Approximately 1% of the genes in eukaryotic genomes code for helicases, and to date, over 100 RNA and DNA helicases have been discovered [2, 3]. These motor proteins function by coupling ATP hydrolysis to mechanical movement to break the hydrogen bonds between complementary base pairs in dsDNA [2]. One such DNA helicase, Pif1, was first identified in a screen for genes that influence the frequency of mitochondrial DNA recombination in *Saccharomyces cerevisiae* [4]. Since its initial discovery, Pif1 has been characterized as a member of the Superfamily 1B group of helicases, which translocate along single-stranded DNA (ssDNA) in the 5’ to 3’ direction [5]. Unlike some members in this superfamily, Pif1 only binds to DNA, but preferentially unwinds RNA-DNA forked duplexes and structured regions such as G-quadruplexes (G4s) and R-loops [3, 6-10].

Following its initial discovery in yeast mitochondria over 20 year ago, Pif1 has been shown to also localize to the nucleus, where it participates in a myriad of DNA transactions. It functions in Okazaki fragment maturation, wherein it lengthens the flap ahead of DNA polymerase δ, allowing RPA to bind to the displaced flap [11]. Recently, it was determined that the rate of replication on lagging strands containing G4s is delayed in the absence of Pif1, underscoring the need for Pif1 to unwind regions that are difficult to replicate due to the presence of DNA secondary structures [12]. Pif1 also stimulates the activity of polymerase *δ* during break-induced replication through bubble migration [13]. Furthermore, prior to mitosis, Pif1 helps to resolve R-loops and aid in protein displacement from tDNA (tRNA genes) [8], while within the ribosomal DNA (rDNA), it is required for maintaining arrest at the replication fork barrier (RFB) [14]. Additionally, Pif1 acts as a negative regulator of telomerase, both at telomeric ends and sites of double-strand breaks (DSBs) where it prevents telomere addition, allowing for the recruitment of DSB repair factors [15-18]. Given Pif1’s involvement in a plethora of cellular activities, it still remains a mystery how its numerous activities are regulated.

Structurally, Pif1 is divided into three domains: N-terminus, helicase core, and C-terminus. A wealth of research has improved our understanding of the structural motifs and related functions of the helicase domain, but in comparison, little is known about the N- and C-terminal domains of Pif1. Currently, these regions are postulated to be modular accessory domains that may serve as regulatory regions [9, 19, 20]. We hypothesize that the functional significance of these domains may help to maintain specific folds that are necessary for protein function and establish points for post translational modifications (PTMs) [21]. Protein PTMs serve to expand a protein’s functional toolbox by altering cellular localization and impacting protein structure, function, and availability while mediating novel interactions with other proteins or nucleic acids [22]. Acetylation of the ε-amino group on a lysine residue is one such PTM that has been studied in the context of modulating chromatin architecture for many decades [23-25]. However, many non-histone proteins are also modified by acetylation, including replication/repair-associated helicases such as Dna2 [26], BLM [27], and WRN [28].

In addition to reports of acetylation of multiple helicases, many functional interacting partners of Pif1 in the Okazaki fragment maturation pathway, namely, Dna2 [26], FEN1 [29], PCNA [30], and RPA [31, 32] are also modified by lysine acetylation. Since the sequence coding for Pif1 is fairly rich in lysine residues (∼10% of the whole sequence), an amino acid that serves as a good target for many PTMs, we were interested in investigating the acetylation dynamics of Pif1. In the current study, we aimed to specifically explore the lysine acetylation status of Pif1, identify the enzymes that dynamically mediate this modification, and elucidate the impact of acetylation on the protein’s cellular functions and alterations to its biochemical activities. Using Pif1-FLAG overexpression constructs, we compared cellular toxicity in wild-type and acetyltransferase or deacetylase mutant strains. Based on the growth phenotypes, we determined that the acetyltransferase (KAT), NuA4 (Esa1), and its counteracting deacetylase (KDAC), Rpd3, are responsible for regulating Pif1 overexpression toxicity *in vivo*. The <u>Pi</u>f1 <u>N</u>-terminal domain (PiNt) was critical for this toxicity regulation, as N-terminally truncated Pif1 (Pif1ΔN) did not behave in the same manner as the full-length Pif1. Additionally, using *in vitro* Piccolo NuA4 (Esa1)-acetylated recombinant protein, we evaluated the impact of this modification on Pif1’s biochemical functions - DNA unwinding, G4 resolvase activity, DNA binding, and ATPase activity-and observed a stimulation in all four activities in the acetylated form when compared to the unmodified form of the protein. The results of our cellular assays and *in vitro* studies indicate that lysine acetylation serves as an important regulator of Pif1 activity within the cell.

## RESULTS

### *Cellular Acetylation Status Modulates Pif1 Overexpression Toxicity* In Vivo

Acetylation dynamics within the cell are tightly regulated by KATs and counteracting KDACs [33]. These lysine modifiers regulate both histone and non-histone proteins, thereby impacting biochemical activities and cellular processes [34]. Because the lysine-rich sequence of Pif1 makes it a target for modification by acetylation, we were interested in defining the effect of this modification on the functional activities of Pif1 within the cell. To understand the impact of global cellular acetylation on the function of Pif1, we initially sought to alter the acetylation dynamics in the cell by creating either KAT or KDAC mutant strains. It has previously been reported that the overexpression of Pif1 in *S. cerevisiae* is toxic to cell growth [35, 36]. Therefore, we used this overexpression toxicity phenomenon to develop a phenotypic assay to determine the impact of cellular acetylation. Our hypothesis was that altering the cellular levels of acetylation may influence Pif1 interactions with other proteins and/or its own function, thereby altering the overexpression toxicity phenotype. A galactose-inducible overexpression plasmid was used to overexpress Pif1 in wild-type, acetyltransferase mutant, and deacetylase deletion mutant cells. However, the growth kinetics (*i.e.*, length of lag phase, doubling rate, and terminal cell density) of *S. cerevisiae* cells are affected by multiple variables in a somewhat stochastic manner [37]. Thus, it can be difficult to compare growth curves between independent experiments using the same strain, let alone comparing strains with multiple genetic backgrounds.

To overcome these limitations of cell growth analyses, we developed a data analysis method to specifically focus on the effect of Pif1 overexpression regardless of strain background. First, cell growth was monitored by measuring the optical density of liquid cultures at 660 nm (OD_660_) over 48 h, and the mean OD_660_ for each strain was calculated. Then, to determine the effect of Pif1 overexpression on growth, the mean OD_660_ of cells grown in galactose-containing medium was divided by the mean OD_660_ of the same strain grown in glucose-containing medium, which strongly represses the *GAL1/10* promoter. Finally, this galactose/glucose ratio for each Pif1-overexpressing strain was normalized to the same ratio from an empty vector control. The normalized growth value for each genotype is interpreted as the toxic effect of Pif1 overexpression in that specific genetic background. The results of these experiments are shown in Figure 1. Here, we recapitulated the toxicity of Pif1 overexpression reported by others [35, 36], finding an approximately 50-60% reduction in wild-type growth upon overexpression of the helicase (Figure 1A).

**Figure 1:**
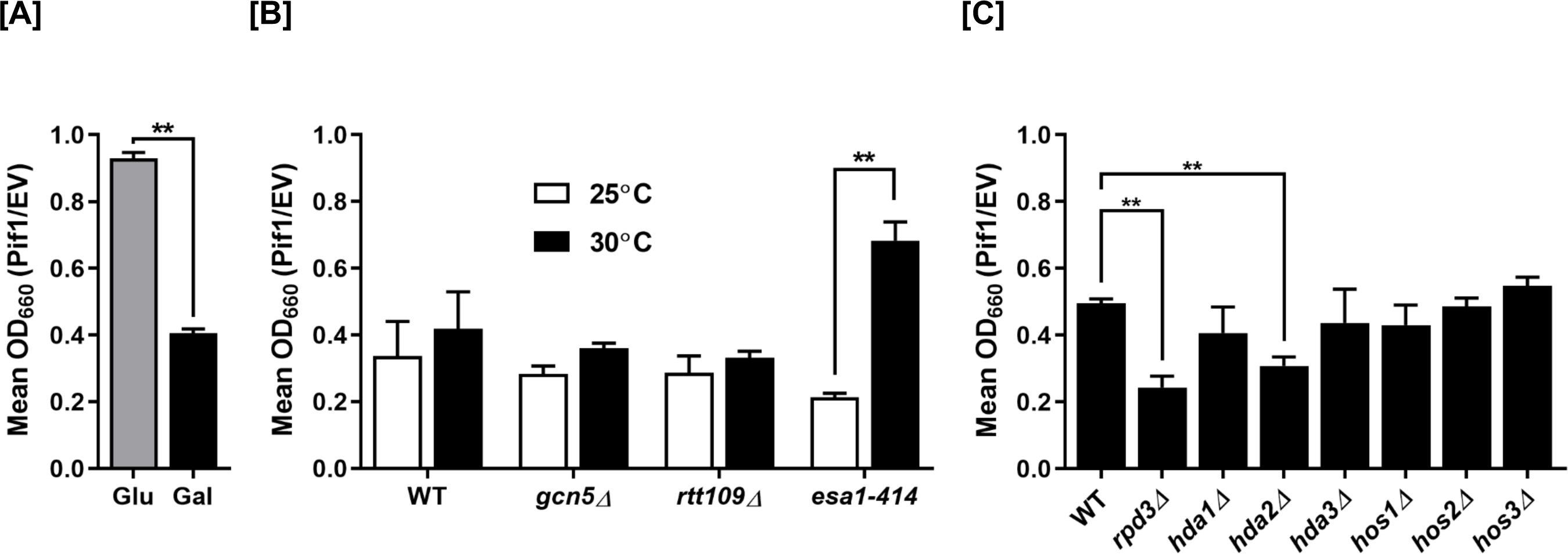
Pif1 over-expression toxicity is altered based on cellular acetylation levels. [A] A galactose-inducible Pif1 expression vector, along with the empty vector, were transformed into wild-type cells. Growth was monitored over 48 h, and the mean OD_660_ of Pif1-expressing strains was normalized to that of the empty vector in both glucose- and galactose-containing media. [B] Pif1 was overexpressed in acetyltransferase deletion (*gcn5Δ* and *rtt109Δ*) or temperature-sensitive (*esa1-414*) mutants at permissive (25°C) and restrictive (30°C) temperatures. [C] Pif1 was overexpressed in the indicated deacetylase mutant strains. The graphed values represent the average of ≥3 independent experiments of technical duplicates, with error bars corresponding to the standard deviation. \*\**p*<0.001.

To determine if acetylation of Pif1 had an effect on overexpression toxicity, we performed similar experiments in KAT mutant cells. Acetyltransferases representative from the Gcn5-related N-acetyltransferase (GNAT) family, the MYST family, and p300/CBP family were chosen for our studies [38]. The genes encoding the Gcn5 (GNAT) and Rtt109 (p300/CBP) KATs are non-essential and can be cleanly deleted, but the catalytic subunit Esa1 of the NuA4 complex (MYST) is essential [39, 40], so the temperature-sensitive *esa1-414* allele [41] was used. This allele has reduced KAT activity at 30°C but full activity at 25°C. Pif1 overexpression in the *esa1-414* background caused significantly (*p* < 0.001) reduced toxicity compared to wild-type when grown at the restrictive temperature (30°C) but not at the permissive temperature (25°C) (Figure 1B). No such effect was observed upon Pif1 overexpression in the *gcn5Δ* and *rtt109Δ* backgrounds, suggesting that NuA4 (Esa1) acetylates Pif1 *in vivo* and that acetylation is connected to Pif1 overexpression toxicity.

To address this hypothesis further, the effects of Pif1 overexpression were assessed in a set of *S. cerevisiae* strains each lacking a single KDAC (Figure 1C). If hypo-acetylation resulted in better growth upon Pif1 overexpression in the *esa1-414* cells, then hyper-acetylation in one or more KDAC-null backgrounds should exacerbate the toxicity. Indeed, we observed significant (*p* < 0.001) increases in Pif1 overexpression toxicity in cells lacking either the KDAC, Rpd3 or Hda2 (Figure 1C). Unfortunately, we could not test for synergistic Pif1 toxicity in a double *rpd3Δ hda2Δ* mutant strain because that combination of KDAC deletions is synthetically lethal (*data not shown*). These results indicate that hyperacetylation of Pif1 in cells lacking Rpd3 or Hda2 leads to increased toxicity upon helicase overexpression. Furthermore, the experiments in Figure 1B suggest that NuA4 (Esa1) mediates Pif1 acetylation *in vivo*, and Rpd3 and/or Hda2 are responsible for deacetylating Pif1 (Figure 1C). It should be noted that NuA4 and Rpd3 are known to have balancing activities *in vivo* [41], lending credence to these results, and thus, we focused on Rpd3 instead of Hda2 herein.

### Pif1 Acetylation is Dynamically Regulated by NuA4 and Rpd3

Because Pif1 overexpression toxicity was significantly altered in specific KAT and KDAC mutant strains, we were interested in directly confirming if the lysine residues on Pif1 were acetylated in these backgrounds. Based on the results obtained in Figure 1B and C, we investigated the acetylation status of overexpressed Pif1-FLAG in the acetylation proficient (*rpd3Δ)* and acetylation deficient (*esa1-414Δ*) strains that displayed significant difference in overall cell viability compared to the wild-type strain. To assess Pif1 acetylation, we immunoprecipitated the lysates with anti-acetyl lysine antibody, followed by immunoblotting with anti-FLAG antibody. Input and phosphoglycerate kinase (Pgk1) served as a loading control in all experiments. The western blot results showed increased Pif1 acetylation in the *rpd3Δ* lysate and decreased Pif1 acetylation in the *esa1-414Δ* lysate compared to the wild-type (compare lane 2 and 3 to lane 1 respectively, Figure 2A), consistent with the results in Figure 1. These results further confirm that Pif1 acetylation is regulated by the action of NuA4 (Esa1) and its counteracting partner, Rpd3.

**Figure 2:**
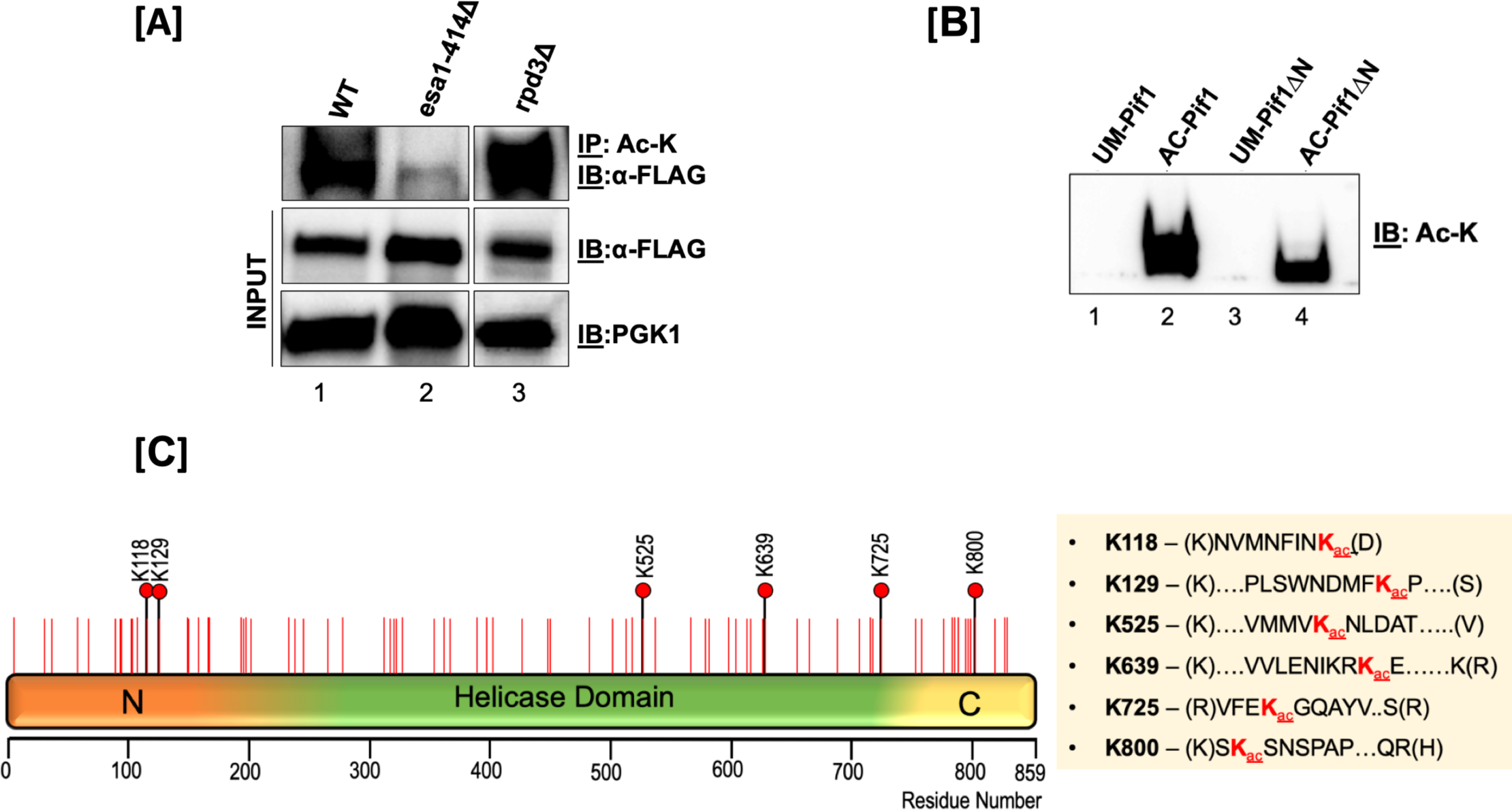
Pif1 is acetylated both *in vivo* and *in vitro*. [A] <u>Top Panel</u>: *S. cerevisiae* lysates from WT (lane 1), *esa1-414* (lane 2), and *rpd3Δ* (lane 3) backgrounds were immunoprecipitated with anti-acetyl lysine (ac-K) antibody-coated Protein G-dynabeads and immunoblotted with anti-FLAG antibody (1:1000); <u>middle Panel</u>: 10% of input immunoblotted with the anti-FLAG antibody and; <u>bottom panel</u>: PGK-1 antibody (1:10,000). [B] Immunoblot of unmodified Pif1 (lane 1), NuA4 (Esa1) *in vitro*-acetylated full-length Pif1 (lane 2), unmodified Pif1ΔN, and NuA4 (Esa1) *in vitro*-acetylated Pif1ΔN probed with anti-Ac-K antibody. [C] Schematic of the full-length Pif1 sequence. The positions of all of the lysine residues in the sequence are denoted with red lines, and all acetylated lysine residues identified by mass spectrometry are denoted with black lines and red filled circles.

To test the efficiency of recombinant Piccolo NuA4 (Esa1/Epl1/Yng2 subunits) for acetylating Pif1, we used a previously established *in vitro* acetylation protocol to modify recombinant Pif1 [29]. Using an anti-acetyl lysine antibody, we were able to detect lysine acetylation of Pif1 on the Piccolo NuA4-modified Pif1 but not on the unmodified form (Figure 2B), confirming that Piccolo NuA4 was capable of acetylating Pif1. Similarly, autoradiography of *in vitro* acetylated Pif1 also showed the helicase to be modified, in addition to the autoacetylation of the Esa1, Epl1, and Yng2 subunits of Piccolo NuA4 (*Supplementary Figure 1*). Further, using tandem mass spectrometry analysis, we were able to identify six acetyl lysine sites on the *in vitro*-modified Pif1: K118, K129, K525, K639, K725, and K800 (Figure 2C). An example of an acetylation spectrum detecting the +42 Da mass shift is shown in *Supplemental Figure 2*. These results establish a Pif1 acetylation signature that defines two sites of modification in the N-terminal domain (K118, K129), three in the helicase domain (K525, K639, K725), and one on the C-terminus (K800). The two modified residues clustered in the PiNt were intriguing because while the function of the PiNt is still being elucidated, it has already been shown to regulate some of Pif1’s activities and alter its overexpression toxicity *in vivo* [36]. To investigate this further, we used an N-terminal domain truncation of Pif1 (Pif1ΔN) lacking amino acids 1-233 to determine the role of the PiNt in Pif1 acetylation. We found that similar to modifying the full-length Pif1, Piccolo NuA4 (Esa1) was also able to *in vitro* acetylate recombinant Pif1ΔN (Figure 2B). Additionally, acetylation of previously determined lysine sites on the helicase and C-terminal domains in the absence of PiNt were confirmed by mass spectrometry.

### The Absence of the PiNt in Acetylation Mutant Strains Impacts Overexpression Toxicity

To assess the contribution of the PiNt to acetylation-dependent toxicity, we used the Pif1ΔN construct in our overexpression toxicity assay, investigating the same genetic backgrounds as in Figure 1. As previously reported [36], Pif1ΔN was less toxic in this assay in wild-type cells than full-length Pif1 (Figure 3A and 3B), exhibiting a toxicity value of ∼0.8 compared to 0.4, respectively. No decrease in toxicity was observed in the *esa1-414Δ* cells nor in any other KAT deletion strain (Figure 3A). Because wild-type cells over-expressing Pif1ΔN already showed robust growth, this result is unsurprising.

**Figure 3:**
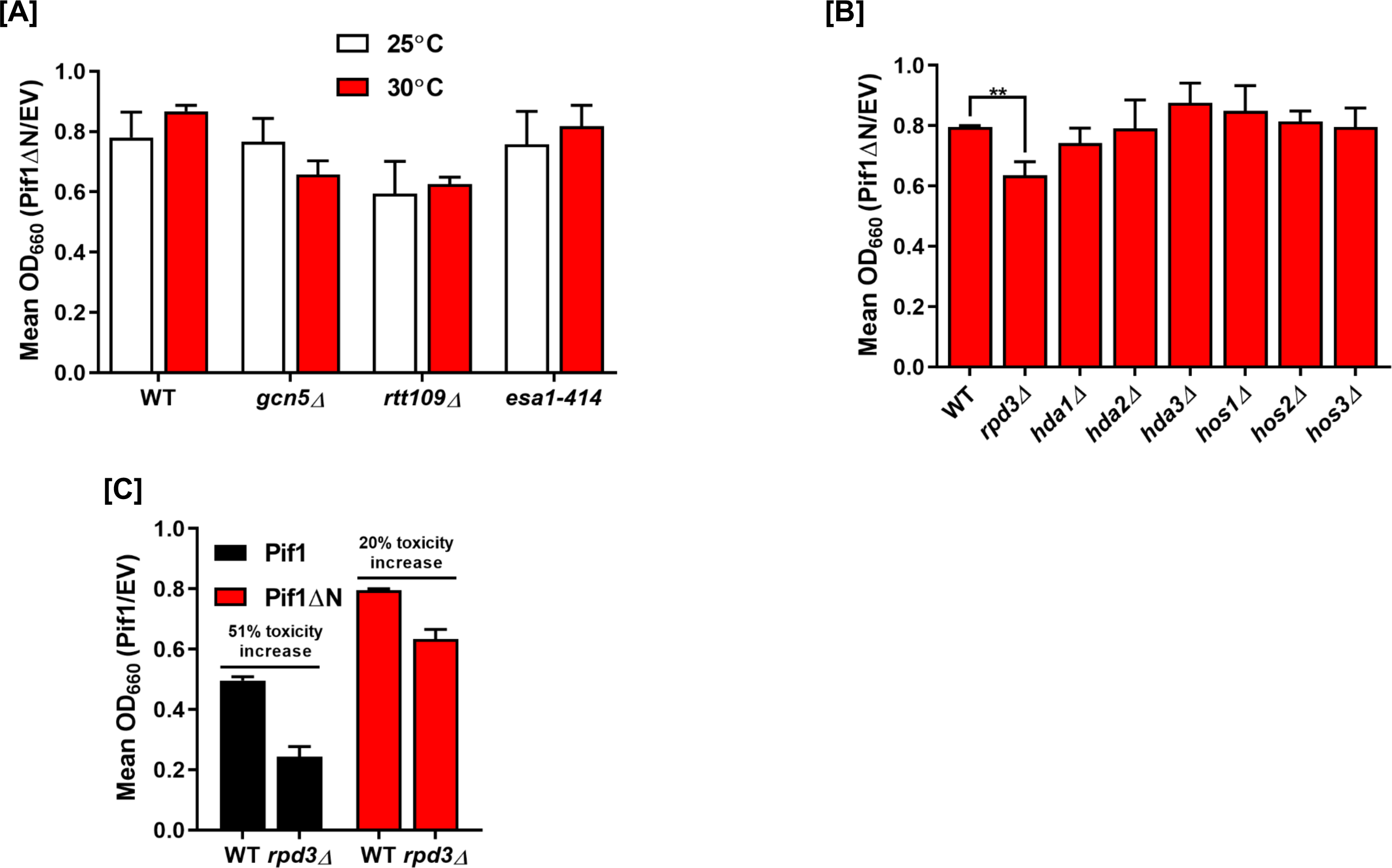
Deletion of the PiNt reduces Pif1’s overexpression toxicity. [A] Pif1ΔN was overexpressed in acetyltransferase mutants at 25°C and 30°C. [B] Pif1ΔN was overexpressed in deacetylase mutant strains. [C] Comparison of the overexpression toxicity of Pif1 and Pif1ΔN in the *rpd3Δ* background. The graphed values represent the average of ≥3 independent experiments of technical duplicates, with error bars corresponding to the standard deviation. \*\**p*<0.01.

Among the KDAC mutant strains, deletion of the PiNt rescued the increased toxicity of Pif1 overexpression in *hda2Δ* cells (Figure 3B). Deletion of *RPD3* still resulted in increased toxicity relative to wild-type cells (*p* < 0.01), upon Pif1ΔN overexpression (Figure 3B), but this still represented a significant growth improvement compared to full-length Pif1 overexpression in *rpd3Δ* cells (*p* < 0.0001). Indeed, growth in *rpd3Δ* cells led to a 51% toxicity increase when Pif1 was overexpressed, compared to only a 20% toxicity increase with Pif1ΔN (Figure 3C). There were no significant effects of Pif1ΔN overexpression in the other KDAC deletion backgrounds. The reduction of acetylation effects on Pif1ΔN toxicity compared to full-length Pif1 indicates that the PiNt is critical for acetylation-altered activity *in vivo*.

### Acetylation Stimulates Pif1’s Helicase Function

Pif1 is a structure-specific helicase, and the order of its preferential unwinding of substrates is RNA:DNA forks > DNA:DNA forks > 5’ tailed DNA:RNA duplex > 5’ tailed DNA:DNA duplex [42]. We designed forked and tailed duplex substrates to aid in determining the impact of lysine acetylation on the helicase activity of Pif1. Because Pif1 is a non-processive helicase, assays were performed using unmodified (UM) and Piccolo NuA4 (Esa1) *in vitro-*acetylated (AC) forms of the protein under multi-turnover conditions, such that more unwound products could be visualized given the experimental parameters used. Under these conditions, Pif1 was able to rebind to its DNA substrate after initial dissociation, allowing for multiple rounds of binding followed by unwinding. The amplitude (*a measure of the percentage of DNA molecules that are completely unwound by the helicase during the course of the reaction)* of unwinding was compared between UM-Pif1 and AC-Pif1 on different cognate helicase substrates. We observed that Pif1 acetylation led to ∼3-fold increase on a DNA-DNA fork (Figure 4A), DNA-DNA tail (Figure 4C), and RNA-DNA tail substrates (Figure 4D) and ∼2-fold increase on an RNA-DNA fork (Figure 4B). Both RNA-DNA substrates had higher amplitudes than their DNA-DNA counterparts, confirming that indeed, the nature of the nucleic acid within the duplex region of the displaced strand dictates Pif1’s preference for certain substrates over others [42]. Furthermore, the Pif1 helicase is additionally known to unwind stable G4 structures, which can hinder DNA replication [12]. Therefore, using a G4 substrate, we determined the impact of acetylation on G4 resolution under similar conditions [43]. Identical to its other preferred substrates, we observed ∼2-fold stimulation of unwinding when Pif1 was acetylated (Figure 4E). Formation of a stable G4 structure was confirmed by performing a synthesis assay using DNA polymerase delta (pol *δ*) in the presence of unmodified and acetylated forms of the helicase (*Supplementary Figure 3*).

**Figure 4:**
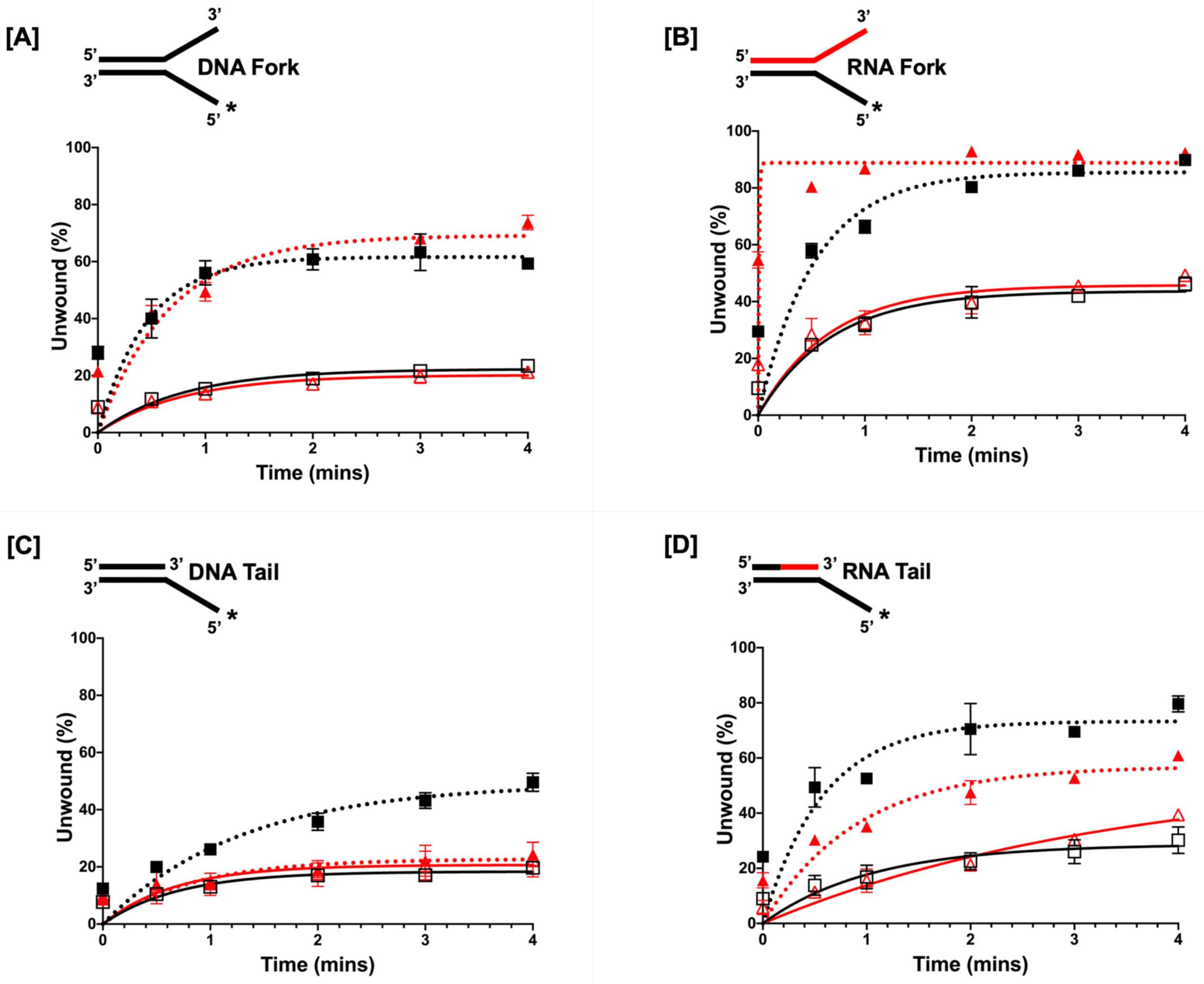
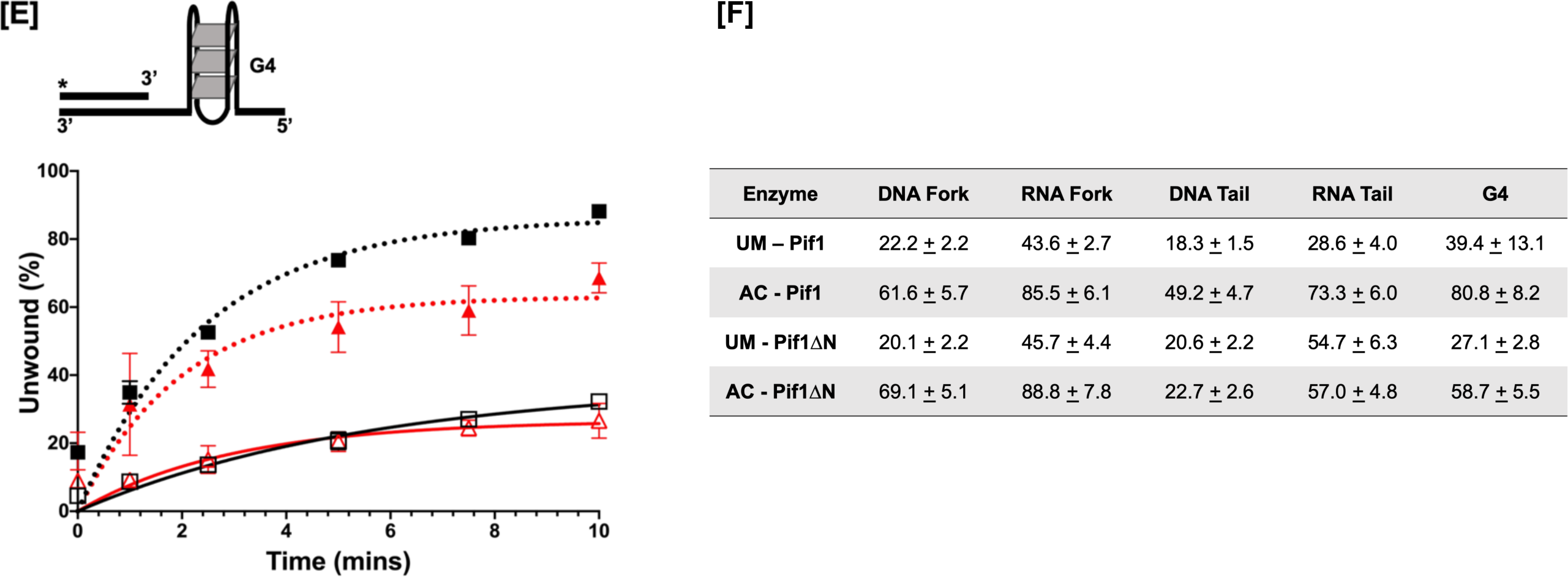
Acetylated Pif1 shows increased helicase activity under multi-turnover conditions. The kinetics of 1 nM full-length Pif1 and Pif1ΔN helicase activity under multi-turnover conditions were assessed over six time points (0’’, 30’’,1’,2’, 3’, and 4’) in the presence of 5 nM IR-labelled [A] DNA fork, [B] RNA fork, [C] DNA tail, and [D] RNA tail substrates. [E] cMyc-G4 template stabilized by K^+^ was annealed to a complimentary 5’-radiolabeled 21-nt ssDNA, and the kinetics of Pif1 structure resolving activity was measured by the accumulation of the radiolabeled substrate over six time points (0”, 1’, 2.5’, 5’, 7.5’, and 10’). In all panels, the black lines with open squares represent UM-Pif1, dashed black lines with filled squares represent AC-Pif1, red lines with open triangles represent UM-Pif1ΔN, and dashed red lines with filled triangles represent AC-Pif1ΔN. F) Comparison of the amplitudes of unwinding by UM- and AC-Pif1 and Pif1ΔN on all substrates. Values are represented as the mean ± standard error of the mean (SEM) of at least three independent experiments.

To study the impact of acetylation on helicase activity in the absence of the PiNt, experiments were also performed using the UM- and AC-Pif1ΔN recombinant protein. Under multi-turnover conditions, acetylation increased the amplitudes of forked duplex unwinding (∼2-3-fold) analogous to the full-length protein (Figures 4A and 4B). However, we observed decreased unwinding of tailed substrates by AC-Pif1ΔN compared to full-length AC-Pif1 (Figures 4C and 4D). These data demonstrate that although Pif1ΔN retains the preference for RNA-DNA substrates, it is not equally affected by acetylation in the same manner as the full-length protein, indicating that the PiNt may be involved in the altered biochemistry of AC-Pif1. Additionally, for the G4 substrate, although acetylation of PifΔN yielded a ∼2-fold increase in the amplitude of unwinding relative to UM-PifΔN, our results suggest that deletion of the PiNt had an impact on Pif1’s interaction with this substrate (Figure 4E). A comparison of the amplitude of unwinding at the highest time-point for every substrate is summarized in Figure 4F. Control experiments showed that neither the presence of Piccolo NuA4 (Esa1) nor acetyl CoA alone impacted helicase activity, but acetylation of Pif1 was necessary to observe stimulation of helicase activity (*Supplementary Figure 4*).

Results from our multi-turnover helicase assays demonstrated that acetylation of Pif1 led to an increase in the amount of unwound products formed compared to UM-Pif1. We then inquired if this increase was due to a change in the rate of Pif1 unwinding when the helicase was modified by lysine acetylation. To address this, helicase assays were performed under single-turnover conditions, where the protein was trapped by the addition of an excess of unlabeled oligonucleotide, permitting only one round of binding and unwinding of its substrate. We ensured that the presence of the protein trap did not affect the unwinding kinetics of the protein (*data not shown*). The rate of unwinding of the DNA-DNA fork by UM-Pif1 was 0.409 ± 0.210 s^-1^, while that of its acetylated form was 0.368 ± 0.132 s^-1^ (Figure 5A). Comparatively, the rate of unwinding of the RNA-DNA fork by UM-Pif1 was 0.344 ± 0.208 s^-1^, and that of AC-Pif1 was 0.344 ± 0.151 s^-1^ (Figure 5B). Thus, the data obtained from the single-turnover assays showed that irrespective of the forked substrate (DNA:DNA or RNA:DNA fork) incubated with the helicase, acetylation did not influence the rate of unwinding. However, we observed that there was still an increase in helicase activity (as measured by the amplitude) on an RNA:DNA fork substrate compared to a DNA:DNA fork substrate, following the previously established phenomenon that Pif1 preferentially unwinds RNA-DNA forks [44]. Additionally, AC-Pif1 displayed ∼3-fold stimulation in the formation of unwound product compared to UM-Pif1 on a DNA:DNA fork substrate, and ∼ 2-fold stimulation on a RNA:DNA fork substrate (Figure 5A and 5B) as measured by their amplitudes. Negligible unwinding of the tailed substrates was found under the experimental conditions used for single-turnover studies (*data not shown*). Interestingly, our results suggest that lysine acetylation does not stimulate Pif1 unwinding by affecting its unwinding rate. Instead, acetylation may make the protein more processive as the amplitude of unwound product formed was increased when substrates were incubated with AC-Pif1. Taken together, these results show that lysine acetylation improves the helicase-catalyzed unwinding of both forked and tailed substrates. They also show that this PTM preserves the protein’s preference and processivity for RNA-DNA hybrids.

**Figure 5:**
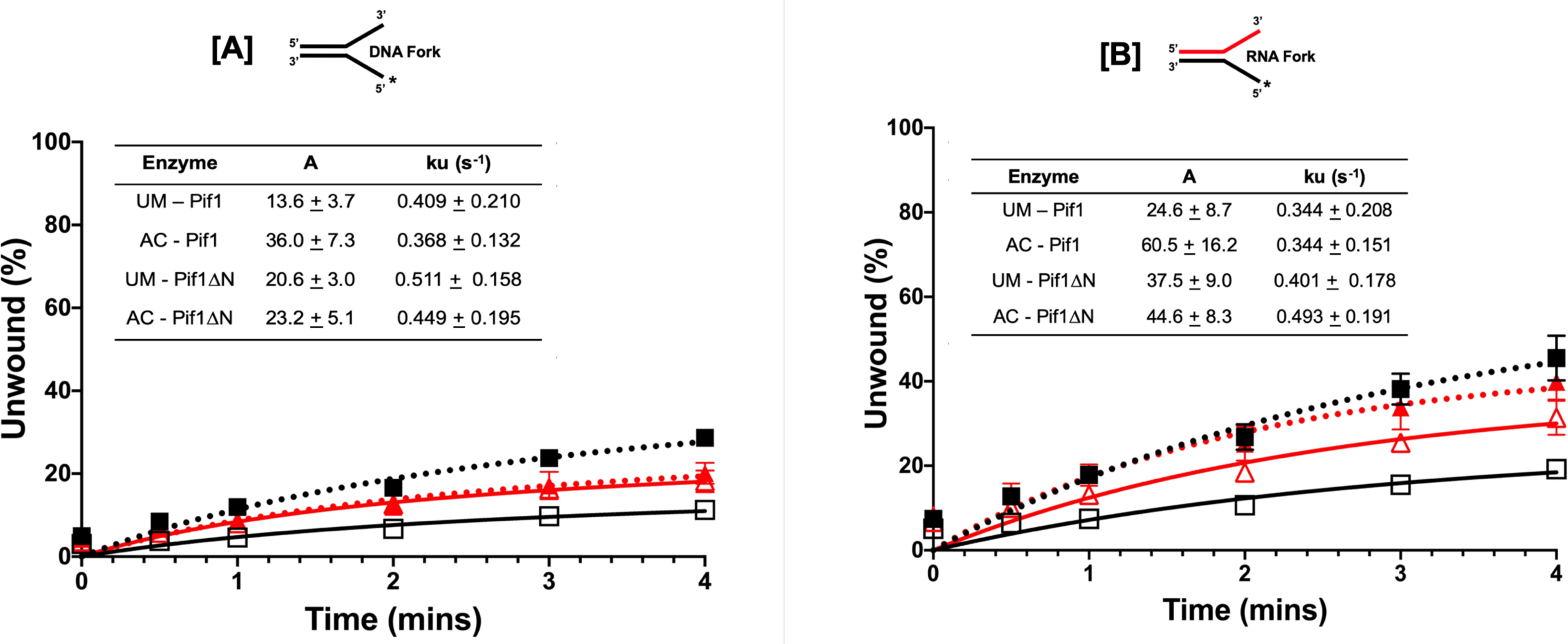
Acetylated Pif1 exhibits increased substrate processivity. The single-turnover kinetics of the unwinding activity of 1 nM full-length Pif1 and Pif1ΔN helicase activity were assessed in the presence of a protein trap (T_50_) and DNA trap (unlabeled 45mer complementary to the displaced strand) over six time points (0’’, 30’’,1’,2’, 3’, and 4’) in the presence of 5 nM IR-labelled [A] DNA fork and [B] RNA fork. Data from the analysis were fit to a first order reaction [A{1-exp[-(ku*X)], in which A is the amplitude of the reaction, ku is the apparent rate constant of unwinding, and X is time. Black lines with open squares represent UM-Pif1, dashed black lines with filled squares represent AC-Pif1, red lines with open triangles represent UM-Pif1ΔN, and dashed red lines with filled triangles represent AC-Pif1ΔN. Values are represented as the mean ± SEM of at least three independent experiments.

### ATPase Activity of Pif1 is Increased upon Lysine Acetylation

As a helicase, Pif1 utilizes the energy produced from ATP hydrolysis to translocate along ssDNA and unwind the DNA duplex. Due to the coupling of its helicase activity to ATP hydrolysis, we sought to determine if acetylation also impacts Pif1’s ATPase activity, using an NADH-coupled spectrophotometric assay [45]. This assay operates on the principle that steady-state hydrolysis of ATP is proportional to NADH oxidation. Because Pif1 is a DNA-stimulated ATPase, we measured the change in absorbance associated with NADH oxidation and ultimately ATP hydrolysis in the presence of a 45-nt ssDNA at 340 nM over a 30-min period. We found a ∼3-fold stimulation in the rate of ATP hydrolysis (Figure 6A) when full-length Pif1 was acetylated compared to its unmodified form, suggesting that this modification co-stimulates the helicase-coupled ATPase activity of Pif1. Comparatively, the rate of ATP hydrolysis of Pif1ΔN displayed no significant difference between the unmodified and acetylated forms of the helicase. Interestingly, we observed that the rate of ATP hydrolysis of PifΔN was higher than that of the full-length protein (Figure 6B), implying that the PiNt may further play a role in regulating Pif1 ATPase activity.

**Figure 6:**
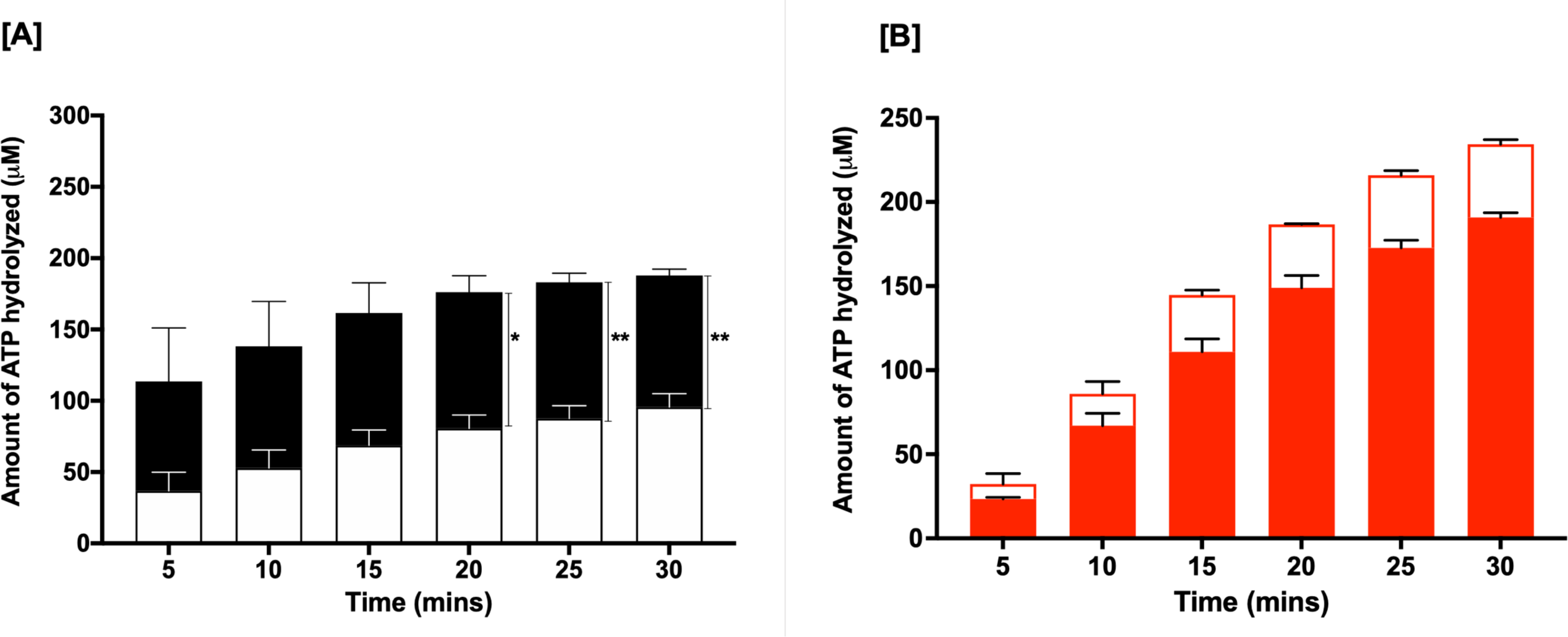
Pif1 ATPase activity is stimulated upon acetylation. Using an NADH-coupled assay, the rate of ATP hydrolysis by [A] UM-Pif1 (open black bars) and AC–Pif1 (filled black bars) and [B] UM-Pif1ΔN (open red bars) and AC-Pif1ΔN (filled red bars) was measured in the presence of 45-nt ssDNA. Values are represented as the mean ± SEM of at least three independent experiments. \**p*<0.05, **p<0.01

### Acetylation Enhances Pif1’s Substrate Binding Ability

This binding to a single-stranded region on the DNA precedes its ATPase activity and ultimately its unwinding function. Therefore, we speculated that the stimulation in helicase-catalyzed unwinding observed upon acetylation might correlate with differential binding of Pif1 to its substrate. To determine this, biolayer interferometry (BLI) technology was used to measure the affinity of UM- and AC-Pif1 to a biotinylated 45-nt ssDNA oligonucleotide immobilized on a streptavidin biosensor. We found that the binding affinity of AC-Pif1 was two-fold stronger than UM-Pif1, with K_D_ values of 7.2 ± 0.3 nM and 19.1 ± 5.1 nM, respectively (Figure 7A). A representative sensorgram showing the binding curves when 125 nM of UM- and AC-Pif1 were incubated with 500 nM of 45 nt ssDNA is presented in Figure 7B. These results show that acetylation of Pif1 leads to higher affinity ssDNA binding. Furthermore, we observed that the UM- and AC-Pif1 ssDNA association rates differed, but their dissociation rates were similar, supporting the hypothesis that the increased amounts of substrate unwound by AC-Pif1 *vs.* UM-Pif1 might be due to faster association and stronger affinity for ssDNA when Pif1 is acetylated. Evaluating DNA:DNA fork binding via EMSAs also revealed a difference in the binding affinities of UM-Pif1 and AC-Pif1, confirming the results obtained from the BLI analyses (Figure 7C).

**Figure 7:**
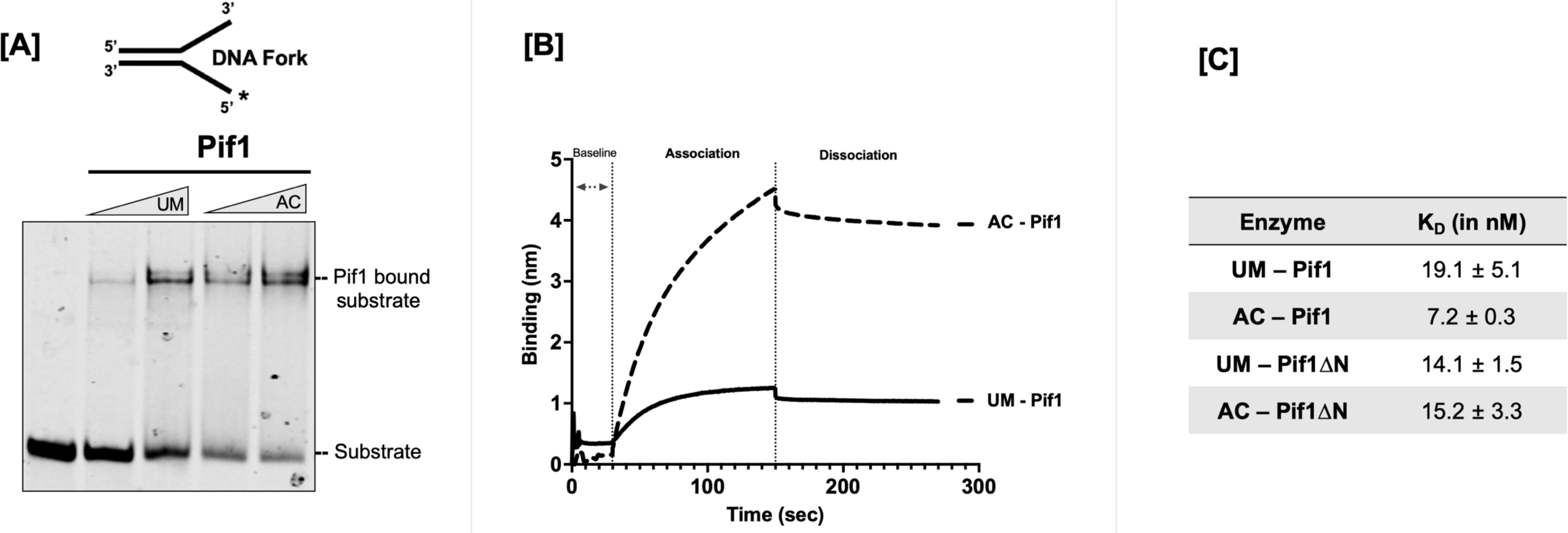
Increased DNA binding affinity is induced by lysine acetylation of Pif1. [A] EMSA of increasing concentrations of Pif1 (100 and 200 nM) in both its unmodified and acetylated form bound to 5 nM DNA fork substrate. [B] Sensogram displaying measured binding kinetics of 125 nM Pif1 incubated with an immobilized biotinylated 45-nt ssDNA using BLItz technology. [C] Binding affinity (K_D_) of the unmodified and acetylated forms of Pif1 and Pif1ΔN.

Based on these results, we hypothesized that AC-Pif1ΔN would show a similar binding trend when compared to UM-Pif1ΔN. However, we observed that acetylation had no impact on the binding affinity of the truncated protein, as the calculated K_D_ values were 14.1 ± 1.5 nM and 15.2 ± 3.3 nM for UM-Pif1ΔN and AC-Pif1ΔN, respectively (Figure 7A). Moreover, these values are similar to unmodified full-length Pif1 (19.1 ± 5.1 nM). This suggests that while Pif1ΔN can bind to ssDNA similar to Pif1, it is not affected by acetylation in the same manner. We hypothesized that acetylation may drive a conformational change in the Pif1 structure that does not occur for Pif1ΔN, and to test this, we next examined changes in Pif1 structure induced by acetylation using limited proteolysis.

### Acetylation Alters the Conformation of Pif1, likely mediated through the N-terminal Domain

To date, there are no published atomic-level structures of full-length *S. cerevisiae* Pif1 including its N-terminal domain [47, 48], presumably due to the challenges presented by the predicted native disorder of the PiNt [36]. Being unable to crystallize full-length UM-Pif1 and AC-Pif1, we instead used limited proteolysis assays to determine if a gross conformational change plays a role in AC-Pif1’s altered biochemical activities. If a conformational change occurs in solution upon Pif1 acetylation, then the protease digestion patterns of UM-Pif1 and AC-Pif1 should differ. We incubated recombinant Pif1 with GluC protease for various lengths of time; GluC was chosen instead of an enzyme such as trypsin or LysC to prevent lysine acetylation from inhibiting the protease. We found that full-length UM-Pif1 was nearly completely degraded within the first 15 min, with lower molecular weight digestion products continuing to form over 60 min (lanes 2 – 4, Figure 8A). In contrast, a proportion of full-length undigested AC-Pif1 remained even after 60 min, with minor small peptide (20-35 kDa) product formation (lanes 6 – 8, Figure 8A).

**Figure 8:**
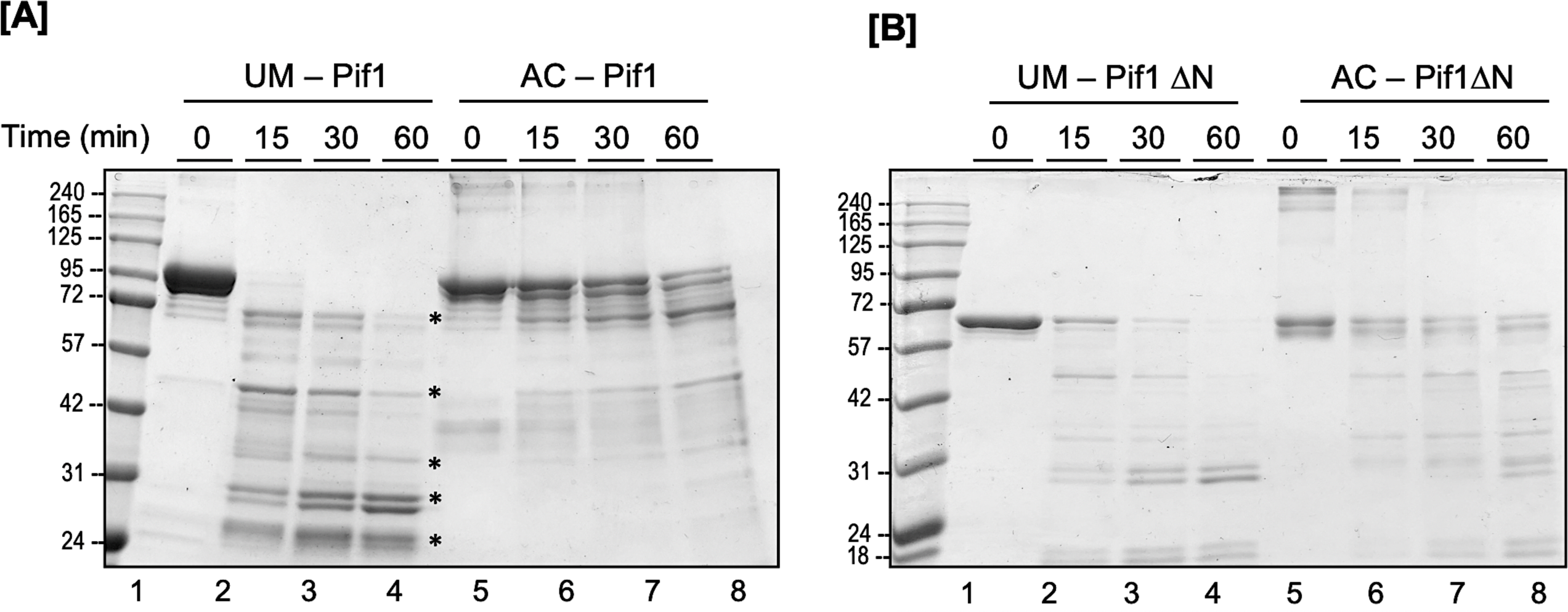
Acetylation of Pif1 induces a conformational change. A time course GluC degradation of unmodified and acetylated [A] Pif1 and [B] Pif1ΔN is shown. The major degradation products are indicated by asterisks on the gel.

Next, we repeated this assay with recombinant UM- and AC-Pif1ΔN proteins to determine the role of the PiNt in acetylation-driven conformational changes. We observed that UM-Pif1ΔN was more resistant to GluC proteolysis than UM-Pif1, with at least a portion of undigested UM-Pif1ΔN evident at all time points (lanes 2-4, Figure 8B). This suggests that the PiNt is a major target of GluC activity in the context of UM-Pif1. Indeed, although we do not know the sequences of the digested species created by limited proteolysis, it should be noted that the highest molecular weight digestion product of UM-Pif1 is approximately the same size as undigested Pif1ΔN (Figures 8A and B), perhaps indicating that the PiNt is easily removed from Pif1 upon GluC digestion. Further, unlike with full-length Pif1, AC-Pif1ΔN was only slightly protected from GluC digestion compared to UM-Pif1ΔN, with both proteins exhibiting similar proteolytic cleavage patterns and rates of digestion (compare lanes 2 – 4 to lanes 6 – 8, Figure 8B). Taken together, these data indicate that acetylation induces a conformational change in Pif1, which occurs either directly in the PiNt or which requires the PiNt for allosteric changes.

## DISCUSSION

While the enzymatic functions of the Pif1 helicase have been extensively characterized, the precise mechanisms by which the activities of this helicase are coordinated to impact a variety of nuclear DNA transactions remain unknown [48-52]. Pif1’s cellular abundance is predicted to be low [53], and aberrant Pif1 levels in the cell lead to deleterious effects. Deletion or depletion of Pif1 from the nucleus results in telomere hyperextension and telomere addition to double-stranded breaks [54]. Conversely, as described previously [35, 36] and in Figure 1A, overexpression of Pif1 is toxic to cells, inhibiting cell growth. These studies demonstrate that the activity of the protein is regulated in the cell by one or more means. Indeed, phosphorylation is known to regulate Pif1’s role in telomere maintenance [55]. However, the role of other PTMs in regulating Pif1 activities has not yet been elucidated. In our current study, we determined the acetylation dynamics of Pif1 *in vivo* and, using *in vitro* biochemical assays, defined alterations to its various enzymatic activities upon modification. We speculate that lysine acetylation is a mechanism used by the cell to regulate the function of Pif1 for different nuclear DNA transactions.

Acetylation of histone tails helps to neutralize the positive charge on lysine residues, causing the destabilization of the chromatin architecture and thereby allowing biological machineries to gain access to the DNA [56]. Enzymes responsible for dynamically modifying histone residues can also interact and acetylate non-histone proteins, including proteins associated with DNA replication and repair [34, 57]. Our toxicity assay for Pif1 overexpression in different KAT and KDAC mutant strains pointed to the KAT, NuA4, and its counteracting partner KDAC, Rpd3, as responsible for cellular Pif1 acetylation (Figures 1B, 1C and 2A). The KAT activity of Esa1 is linked to cell cycle progression, potentially by regulating transcription [58]. In *S. cerevisiae*, Rpd3 is also associated with cell cycle control by regulating replication origin firing [59]. Both Esa1 and Rpd3 also play an important role in the DNA repair process, albeit these studies are in connection with histone acetylation [60]. Because both the acetylation modifiers are in close contact with chromatin during cell cycle progression and repair, it is not surprising that they would also play a role in modifying Pif1, a helicase associated with replication and repair. Our Pif1 toxicity assay, while quantitative, will not catch subtle modifications (singular or multiple lysine acetylation on Pif1) that other redundant KATs and KDACs may be able to accomplish. However, we also found Hda2 to play a subtle role in the deacetylation process, though, its impact may not have been as high as Rpd3, based on the toxicity studies (Figure 1B). Thus, although NuA4 (Esa1) and Rpd3 may serve as the primary modifiers of Pif1, we cannot rule out the activity of other KATs and KDACs in regulating Pif1 acetylation, because these modifiers display redundancy in their cellular functions [61]. Although we did not evaluate if acetylation of Pif1 is coordinated along with cell cycle phases in our current study, previous work demonstrates that acetylation of other replication proteins, including FEN1 and PCNA, does not display cell cycle specificity [62, 63]. Nonetheless, considering that Pif1 partakes in multiple DNA transactions, lysine acetylation may be specifically regulated in response to a genome maintenance event requiring alterations to specific activities of Pif1.

Acetylation of recombinant proteins has its own caveats, including that the *in vitro* acetylation reaction never goes to completion and, in the absence of other protein regulators, tends to be promiscuous [64]. However, the six lysine residues we report to be modified on Pif1 were acetylated in multiple independent *in vitro* reactions, thus making them robust potential targets for modification by Piccolo NuA4 (Esa1) (Figure 2C). Of these lysine residues, two resided in the PiNt, three in the helicase domain, and two in the C-terminus. The fact that this modification is not limited to a certain segment/domain of the protein suggests that many of the various biochemical properties of the protein could be impacted. Pif1’s helicase core alone houses the seven conserved amino acid motifs common to this family where the direction of ssDNA translocation is determined, ATP hydrolysis, and ssDNA binding occur [65-67]. Lysine acetylation of the BLM helicase is similarly spread across its different domains, allowing for regulation of its functions during DNA replication and the DNA damage response [27].

Helicase assays performed using *in vitro*-modified Pif1 revealed a significant stimulation in its unwinding activity upon acetylation compared to the unmodified form. This stimulation was apparent on all substrates tested, even though the levels of stimulation differed based on the specific substrate being unwound (Figure 4F). The single-turnover and multi-turnover helicase reactions confirmed that the increased helicase unwinding was due to increased processivity and not due to a faster rate of unwinding (Figures 4 and 5). In addition to Pif1’s helicase activity, its ATPase function was also stimulated when the protein was acetylated (Figure 6). However, because Pif1 is a DNA-stimulated ATPase, it is difficult to determine if this stimulation is due to faster ATP hydrolysis or if faster DNA binding allows ATP hydrolysis to occur more rapidly. Future work using order-of-addition ATPase assays could help to delineate between these two possibilities.

Characterization of DNA binding activity demonstrated increased binding by AC-Pif1 compared to UM-Pif1 (Figure 7). At a first glance, this observation is counterintuitive because one would expect lower nucleic acid binding affinity when the positive charge on lysine is neutralized by acetylation. However, Pif1 was acetylated at a single site, K725, located within the DNA binding domain (DBD) [68], while the other modified sites were found throughout the protein. Alterations in binding activities could depend on sites modified within and outside the DBD, and how each of those individual lysine charge neutralization events impact the overall binding of a protein. Of the six Pif1 lysine residues we found to be acetylated *in vitro*, one residue, K525, is conserved in hPIF1 (K485). K485 makes contact with ssDNA, and mutation of this residue to alanine results in decreased ssDNA binding affinity [49]. In yeast, K525 may be important for the regulation of Pif1 DNA binding and acetylation-altered activity, which mutational analysis would elucidate further. Acetylation of other proteins, such as p53 [69], Gata-1 [70], and Stat3 [71], all serve as examples of proteins displaying increased binding affinities for specific DNA substrates when acetylated. Interestingly, acetylation of replication protein A (RPA) promotes its displacement from ssDNA during DSB repair [72], and acetylated FEN1 displays lower substrate binding affinity [63]. All of these proteins are hypothesized to undergo conformational changes upon acetylation, which may alter their ability to interact with and bind to their cognate substrates.

Pif1 has a large N-terminal domain (PiNt) making up almost one-third of the protein, and this domain is predicted to be natively disordered [36]. Due to the ability to mutate the N-terminus and still retain helicase activity, we focused on characterizing the two acetylation sites on the N-terminal domain. Overexpression toxicity assays revealed that deletion of the PiNt resulted in lesser toxicity compared to wild-type Pif1 when cellular acetylation dynamics were altered (Figure 3). Because the PiNt is predicted to be natively disordered, we hypothesized that it might undergo a conformational change when Pif1 is acetylated, thus impacting the biochemical properties of the helicase domain. Acetylation has been documented to cause conformational changes in a number of other proteins. For instance, PCNA is acetylated in response to DNA damage, and this induces long-range conformational changes in the protein, distal from the acetylation site [62]. Similarly, the DNA binding protein TCF4 is suggested to change conformations when acetylated in a complex with DNA [73], and Beta 2-glycoprotein changes from a closed to open conformation upon acetylation [74].

Limited proteolysis is a method of detecting protein conformational changes that does not require crystallization or large amounts of protein, which are currently both obstacles when working with AC-Pif1. The altered digestion and degradation patterns in the acetylated form of Pif1 compared to the unmodified form indicate changes in the tertiary structure of the protein (Figure 8). Similarly, because AC-Pif1ΔN displayed the same digestion pattern as UM-Pif1ΔN, we speculate that the PiNt is necessary for the acetylation-based conformational change. It may be that the acetylated residues in the helicase and/or C-terminal domain are responsible for changes in the PiNt, similar to the allosteric changes that acetylation drives in PCNA. Alternatively, a combination of residues in every domain might require acetylation for these changes to take place. The structure of the PiNt is unknown, but the transition from a closed to an open conformation like Beta 2-glycoprotein upon acetylation could explain how Pif1 ssDNA binding affinity increases. Additional study, including high-resolution structures of full-length UM-Pif1 and AC-Pif1, is needed to address these questions and further understand how acetylation affects Pif1 structure.

The impetus for studying Pif1 lysine acetylation was triggered by the observation that multiple proteins involved in the Okazaki fragment maturation pathway are also acetylated. Studies *in vitro* have shown that Pif1 promotes increased strand displacement synthesis by the lagging strand DNA pol *δ* [75]. Increased strand displacement allows for the creation and cleavage of longer 5’ displaced flaps. While genetic and biochemical studies support a redundant alternate long flap pathway for Okazaki fragment maturation, this model is largely based on *in vitro* reconstitution assays. One hypothesis for creating longer flaps in the cell is to completely remove the initiator RNA/DNA primer on the lagging strand that is synthesized by the error-prone DNA polymerase α [76]. Another possibility to consider is if Rad27^FEN1^ disengages from the replisome, it may unintentionally allow for the creation of longer flaps within the cell, thereby necessitating an alternate pathway for flap processing. Along with acetylation of other Okazaki fragment proteins, including Rad27^FEN1^ and Dna2, Pif1 modification may push the creation of longer flaps during the maturation process, allowing for higher fidelity synthesis (Figure 9). Recent evidence from the Rass group provides an alternate explanation for the observation that the lethality of *dna2Δ* is suppressed by *pif1Δ* [77]. In this study, they show that in *dna2Δ* cells, Pif1 mediates checkpoint activation following replication stress, which leads to replication fork stalling. These stalled replication forks are resolved through break-induced replication or recombination-dependent replication (RDR), both of which utilize the Pif1 helicase for efficient D-loop structure resolution. Acetylation of Pif1 may also play a role in coordinating the checkpoint response and recruitment of proteins during stalled replication.

**Figure 9:**
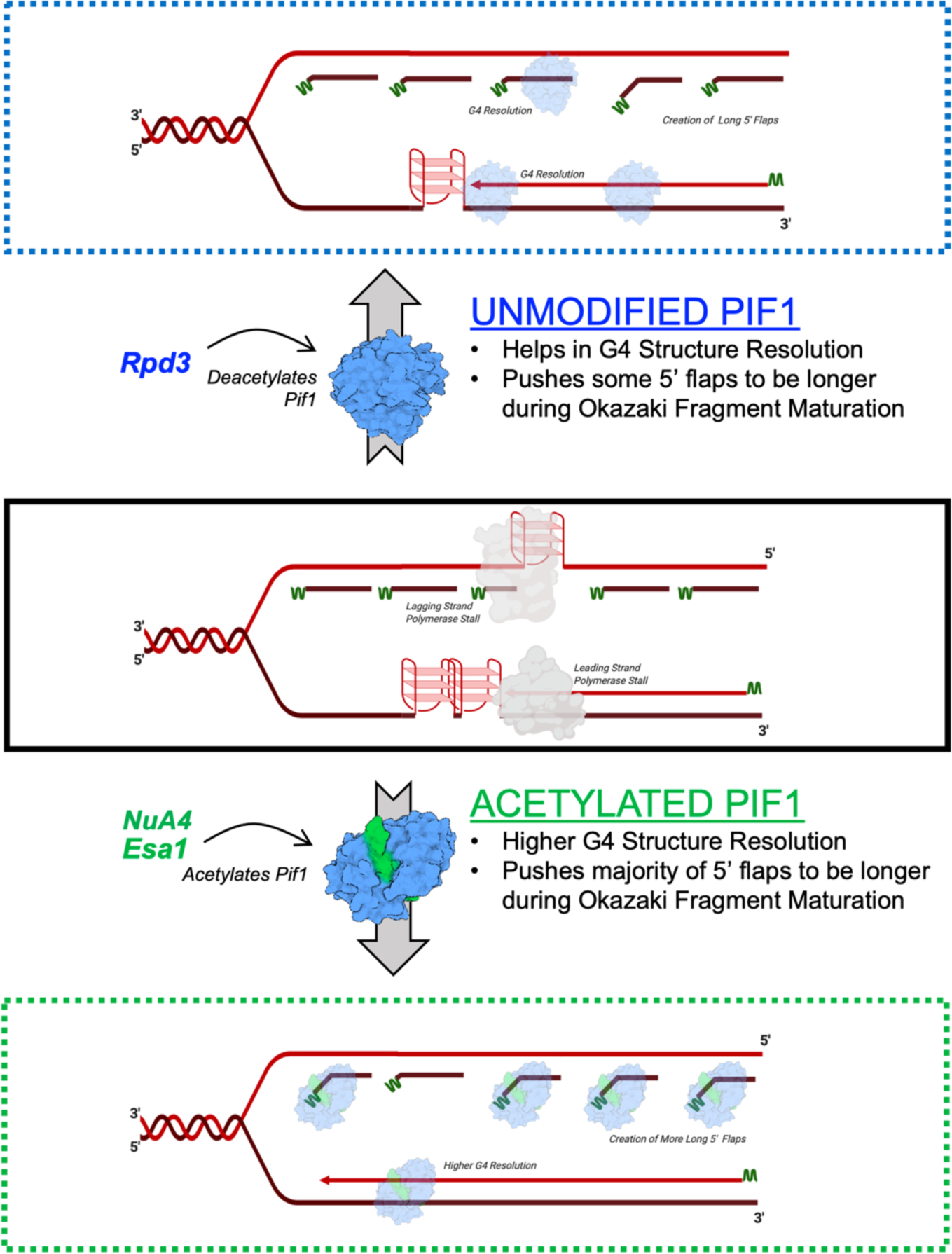
Model for regulation of Pif1 activities by acetylation. <u>Middle Panel</u>: Replication fork with two tandem G4 structures on the leading strand, one G4 structure on the lagging strand (replication stalled), and Okazaki fragments (RNA primers indicated in green). <u>Top Panel</u>: UM-Pif1 (or deacetylated by the KDAC Rpd3) is able to resolve one G4 structure on the leading strand and the G4 structure on the lagging strand, allowing for synthesis of Okazaki fragments. The presence of UM-Pif1 (or deacetylated Pif1) allows for the displacement of a few long 5’ flaps during Okazaki fragment maturation. <u>Bottom Panel</u>: AC-Pif1 (by the KAT,NuA4 (Esa1)) resolves the tandem G4 structures on the leading strand, resolves and allows synthesis over the G4 structure on the lagging strand, and pushes the majority of the flaps to be longer during Okazaki fragment maturation.

AC-Pif1 was also shown to be more efficient at G4 structure resolution (Figure 4E). Though the unmodified form of Pif1 is capable of efficient G4 resolution, due to increased binding affinity of the acetylated form, AC-Pif1 could potentially resolve tandem G4 structures more effectively than UM-Pif1 (Figure 9). Thus, while finely regulated concentrations of AC-Pif1 may be critical to the maintenance of overall genome stability, increases in AC-Pif1 levels may overwhelm the replication machinery by generating longer ssDNA segments, leading to cellular toxicity. Other studies have shown similar results supporting this hypothesis. For instance, SV40 T-antigen inhibits Okazaki fragment processing when in a higher-concentration hexameric state, but it supports Okazaki fragment processing when in a lower-concentration monomeric state [78]. Dysregulation of Rad5, an enzyme involved in post-replication repair in yeast and humans, leads to cisplatin sensitivity when both deleted or overexpressed [79]. The study of human PIF1 (hPIF1) has also demonstrated the deleterious consequences of Pif1 helicase misregulation. The transfection of cultured primary neurons with hPIF1 is toxic, increasing the risk of cell death [80]. Further, human tumor cells rely on hPIF1 for protection from apoptosis [81], whereas hPIF1 depletion in normal cells increases replication fork arrest [82]. The L319P hPIF1 mutant is linked to breast cancer and cannot suppress the lethality of Pfh1 deletion in *S. pombe* [83], indicating that mutant hPIF1 activity could also lead to cancer cell growth. These studies demonstrate that regulation of PIF1 family helicases is critical not just in *S. cerevisiae* but also in humans, and as such, acetylation of hPIFif1 may be a conserved modification used by the cell to regulate hPIF1 activity.

In conclusion, we propose that lysine acetylation of Pif1 is a regulatory mechanism that dynamically alters the cellular enzymatic activity of the helicase. While this study serves as the first report of acetylation-based regulation of Pif1, many questions remain unanswered. The precise timing and cellular triggers of Pif1 acetylation are still unknown. Likewise, how acetylation affects Pif1’s interactions with other proteins and cellular localization remain to be determined. Here, we report the importance of the N-terminus in regulating acetylation-dependent Pif1 activity. While the C-terminus, also predicted to be disordered, serves as an important point of contact for PCNA during BIR [75], we are currently unaware of how specific lysine residue acetylation impacts this domain. Crosstalk between PTMs is commonplace, and as such, acetylation may be connected to Pif1 phosphorylation or other lysine residue-dependent PTMs. Studies designed to answer these and other remaining questions regarding Pif1 activity and regulation will be important to further our understanding of how Pif1 achieves its multi-faceted role of maintaining genomic integrity.

## EXPERIMENTAL PROCEDURES

### Strains, media, and reagents

The *S. cerevisiae* strains used are listed in Supplementary Tables 1 and 2. The cells were maintained on rich medium (YPD) or synthetic drop-out medium and transformed with overexpression plasmids using standard methods. *Escherichia coli* strain NiCo21(DE3) (New England Biolabs) was transformed with the pLysS plasmid (Novagen) to create the NiCo21(DE3) pLysS strain. The *E. coli* cells were maintained on LB medium supplemented with antibiotics (50 μg/ml kanamycin, 34 μg/ml chloramphenicol, and/or 100 μg/ml ampicillin as needed). Liquid cultures were grown in 2× YT medium for protein overproduction and supplemented with the same antibiotics. dNTPs were purchased from New England Biolabs (Ipswich, MA). Oligonucleotides were purchased from IDT (Coralville, IA) and are listed in Supplementary Table 4. Chemical reagents were purchased from Thermo-Fisher, Sigma, or DOT Scientific.

### Overexpression toxicity assays

Plasmid pESC-URA was used for the galactose-induced overexpression of proteins in *S. cerevisiae*. Empty pESC-URA vector or pESC-URA-Pif1 (WT or mutant) was transformed into the indicated yeast strains, and transformants containing the plasmid were selected on SC-Ura drop-out media. Fresh transformants were then grown in liquid SC-Ura medium containing 2% raffinose for 16 h, the cells were harvested and washed with sterile water, and then diluted to an OD_660_ of 0.01 in SC-Ura supplemented with either 2% glucose or galactose. A 200-µL volume of each culture was added in duplicate to wells in 96-well round bottom plates, and each well was overlaid with 50 µL of mineral oil to prevent evaporation. The plate was monitored using a Synergy H1 microplate reader (BioTek), taking OD_660_ measurements at 15-min intervals for 48 h, with linear shaking occurring between readings. The plate reader also incubated the cells at 25 or 30°C as indicated. The mean of the OD_660_ readings for each Pif1-expressing strain grown in galactose was divided by the mean OD_660_ of the same strain grown in glucose. This mean value was normalized to that of cells from the same genetic background containing empty vector to produce a toxicity value for each Pif1 variant in each yeast genotype. Plasmids used in this study are detailed in Supplementary Table 3.

### Protein purification

*S. cerevisiae* Pif1 was over-expressed in NiCo21(DE3) pLysS cells and purified as previously reported [18], with slight modification. To increase the yield of SUMO-Pif1, up to 10 mL TALON resin was used in the form of tandem 5-mL TALON HiTrap columns in the initial capture of recombinant protein from lysate. The Pif1ΔN mutant [9] lacked the first 233 amino acids of the helicase. Recombinant Pif1ΔN protein was expressed and purified in an identical manner to full-length wild-type (WT) Pif1. The *S. cerevisiae* Piccolo NuA4 complex (consisting of Esa1, Epl1, and Yng2) was over-expressed using the polycistronic expression system developed in the Tan laboratory and purified from BL21(DE3) pLysS cells as previously described [84]. *S. cerevisiae* pol δ was purified by co-expressing the pol 3, pol 31, and GST-pol 32 plasmids in BL21(DE3) cells and purifying as previously described [85].

### *In vitro* acetylation

*In vitro* acetylation of Pif1 was performed using two complementary methods. In the first method, purified recombinant Pif1 (Pif1 or Pif1ΔN) was incubated with the Piccolo NuA4 complex (Esa1/Epl1/Yng2) in the presence of acetyl-CoA in 1X HAT buffer (50 mM Tris-HCl (pH 8.0), 10% (v/v) glycerol, 150 mM NaCl, 1 mM dithiothreitol (DTT), 1 mM phenylmethylsulfonyl fluoride, and 10 mM sodium butyrate) for 30 min at 30°C. For proteins analyzed via mass spectrometry, DTT was omitted from the reaction buffer.

In the second method, Pif1 was *in vitro* acetylated using the same protocol as above, but the reactions were incubated for 60 min. Subsequently, the reaction mixture was loaded onto a TALON affinity column. Because Esa1 was 6X-His tagged, the Piccolo complex remained bound to the column, whereas the acetylated Pif1 was eluted by the column wash buffer (50 mM sodium phosphate (pH 7.5), 300 mM NaCl, 1 mM PMSF, 5 mM *β*-mercaptoethanol, 10% (v/v) glycerol, and 7 µg/uL pepstatin A). The eluate was analyzed by SDS-PAGE and Coomassie staining and found to contain no contaminating proteins or subunits from the acetyltransferase.

A ratio of 1:1:10 [Pif1/ Pif1ΔN):acetyltransferase:acetyl-CoA] was maintained for all acetylation reactions.

Results described in this report used AC-Pif1 that was obtained using the second method. However, results for all biochemical assays were confirmed using AC-Pif1 obtained using both methods.

### Mass spectrometry

Tandem mass spectra from *in vitro* Piccolo NuA4 (Esa1)-acetylated full-length Pif1 were collected in a data-dependent manner with an LTQ-Orbitrap Velos mass spectrometer running XCalibur 2.2 SP1 using a top-fifteen MS/MS method, a dynamic repeat count of one, and a repeat duration of 30 s. Enzyme specificity was set to trypsin (or Lys-C when cleaved with this protease), with up to two missed cleavages permitted. High-scoring peptide identifications were those with cross-correlation (Xcorr) values ≥1.5, delta CN values ≥0.10, and precursor accuracy measurements within ±3 ppm in at least one injection. A mass accuracy of ±10 ppm was used for precursor ions, and a mass accuracy of 0.8 Da was used for product ions. Carboxamidomethyl cysteine was specified as a fixed modification, with oxidized methionine and acetylation of lysine residues allowed for dynamic modifications. Acetylated peptides were classified according to gene ontology (GO) annotations by Uniprot. Lysine residues identified as being modified in three or more independent *in vitro* reactions, cleaved with either trypsin or Lys-C, are reported.

### Western blotting

Pif1 protein and the acetylation levels of over-expressed Pif1-FLAG were probed in WT, *rpd3Δ*, and *esa1-414* cells. Cells were grown in YPD or selective media and harvested at OD_660_ = 1.0. Harvested cells were lysed by incubation for 10 min at 95°C in yeast lysis buffer (0.1 M NaOH, 0.05 M EDTA, 2% SDS, and 2% β-mercaptoethanol). Then, 5 µL of 4 M acetic acid was added for every 200 µL lysate, and the mixture was vortexed for 30 s. Lysates were incubated again at 95°C for 10 min, and the soluble fraction was collected by centrifugation [86]. Overexpression levels of Pif1 and Pif1ΔN were detected by immunoblotting with anti-FLAG antibody (Millipore Sigma A8592). For loading controls, the levels of Pgk1 were detected by immunoblotting with an anti-Pgk1 antibody (Fisher 22C5D8). Acetylation of cellular Pif1 was detected by immunoprecipitating cell lysate with anti-acetyl lysine antibody (CST 9441) resin and immunoblotting with an anti-FLAG M2 antibody (Millipore Sigma A8592). Specifically, Protein G Dynabeads (Invitrogen 10007D) were incubated with anti-acetyl lysine antibody (CST 9441) with end-over-end rotation for 4 h at 4°C, followed by the addition of 1 mg of cell lysate, which was then rotated overnight at 4°C. The beads were washed three times in the washing buffer provided with the Dynabead kit, and the beads were resuspended in 2X Laemmli buffer, boiled, and analyzed by SDS-PAGE followed by western blotting using the anti-FLAG M2 antibody. The level of acetylation was determined by comparing expression of Pif1 to the amount of acetylated Pif1.

To detect *in vitro*-acetylated Pif1, 2.5 μM of purified protein, unmodified (Um-Pif1) or acetylated (Ac-Pif1; acetylated protein obtained by the second method above), was separated by 4-15% gradient SDS-PAGE and immunoblotted with the anti-acetyl lysine antibody.

### Oligonucleotides

Synthetic oligonucleotides were purchased from Integrated DNA Technologies. The lengths and sequences (5’-3’) of each oligonucleotide are listed in Supplementary Table 4. For helicase assays, the template was 5’-labeled using the IR 700 dye synthesized by IDT. The IR-labelled template primer (T1) was annealed in IDT duplex buffer to oligonucleotide D1, D2, D3, or D4 to generate the DNA fork, RNA fork, DNA tail, or RNA-DNA tail substrate (respectively) in a 1:4 ratio. Oligonucleotides employed in BLItz assays contained a 3’ biotin tag to allow for binding to the streptavidin biosensors. The G4 template (T2) was radiolabeled with [γ-^32^P] ATP from Perkin Elmer and incorporated at the 5′ end using polynucleotide kinase as previously described [29]. The template was further purified on a 12% sequencing gel containing 7 M urea. The radiolabeled T2 oligonucleotide was annealed to oligonucleotide D5 in a buffer containing 20 mM Tris (pH 8.0), 8 mM MgCl_2_, and 150 mm KCl in a 1:4 ratio. All annealing reactions were incubated at 95°C for 5 min and then slowly cooled to room temperature as previously described [43].

### Electrophoretic mobility gel shift assay (EMSA)

The binding affinities of UM-Pif1 and AC-Pif1 were measured by incubating increasing concentrations of the protein (100 and 200 nM) with 5 nM DNA fork substrate in 1X EMSA buffer (50 mM Tris-HCl (pH 8.0), 2 mM DTT, 30 mM NaCl, 0.1 mg/mL bovine serum albumin, and 5% (v/v) glycerol). The reactions were incubated at 30°C for 10 min, and samples were loaded onto a pre-run 8% polyacrylamide native gel. The gel was electrophoresed at a constant 250 V for 1 h and imaged using an Odyssey imaging system (700-nM filter). Using Image Studio, the densitometry of each band was used to calculate binding affinity with the equation: *[(bound product) / (bound product + substrate remaining) *100]*.

### BLItz analysis

To measure the binding kinetics of the different forms of Pif1 to single-stranded (ss)DNA, 500 nM of a biotinylated 45-nt substrate (T1) was diluted in 1X HAT buffer and coated onto a streptavidin dip read biosensor for 120 s (Pal Forte Bio, CA, USA.) A baseline was established for all biosensor reads by immersing them in 1X HAT buffer for 30 s. Following the baseline reading, 4 μL of the corresponding form of Pif1 at varying concentrations (62.5, 125, 250, and 500 nM) was applied to the biosensor for 150 s to measure association. Upon completion, the biosensor was then immersed in 550 μL of 1X HAT buffer to establish dissociation kinetics for another 150 s. The shift in wavelength was recorded, and the binding affinity (K_D_) was analyzed using ForteBio software.

### Helicase assay

The unwinding efficiency of the different forms of Pif1 was assessed on a wide variety of substrates using either multi- or single-turnover assays. For multi-turnover reactions, 5 nM substrate was incubated with 1 nM Pif1 at 30°C for 0, 0.5, 1, 2, 3, or 4 min. Reactions were performed in helicase buffer (50 mM Tris-HCl (pH 8.0), 2 mM DTT, 30 mM NaCl, 0.1 mg/mL bovine serum albumin, 5% (v/v) glycerol, 4 mM MgCl_2_, and 8 mM ATP) and terminated with 80 mM EDTA, 0.08% SDS, and 50% formamide (final concentration) as previously described [43]. For single-turnover reactions, 1 μM protein trap (T50) and 75 nM cold trap complementary to the labelled strand were added to prevent reannealing. Reactions under these conditions were started by adding 8 mM ATP and 4 mM MgCl_2_. Samples were loaded onto pre-run 8% native polyacrylamide gels and electrophoresed for 30-45 min at 250 V. Gels were imaged using an Odyssey imaging system (700-nM filter) and quantified using Image Studio. The percentage of substrate unwound was calculated using the following equation: *% unwound = [(unwound product) / (unwound product + substrate remaining) *100]*. Unwinding data were fit to the equation *A(t) = A(1-e*^*kut*^) using Graphpad Prism, where A is the amplitude of product formation, k_u_ is the rate of unwinding, and t is time [87].

### ATPase assay

ATP hydrolysis was measured using an NADH coupled assay in the presence of 3 μM ssDNA (unlabeled T1) and 10 nM helicase. The reaction buffer (10 mM ATP, 10 mM MgCl_2_, 1 mM phosphoenolpyruvate, 10 U/mL pyruvate kinase, 16 U/mL lactate dehydrogenase, and 0.8 mM NADH) was pre-loaded into 96-well plates. To start the reaction, protein and DNA were added, and absorbance readings at 340 nm were recorded every 60 s for 30 min using a BioTek Cytation 5™ multi-mode plate reader. The rate of hydrolysis was determined as previously reported [88].

### Proteolysis assay

Proteolysis assays were performed at 30°C for up to 60 min. Pif1 proteins were diluted in 25 mM Na-HEPES, 5% glycerol, 50 mM NaOAc, 150 µM NaCl, 7.5 mM MgOAc, and 0.01% Tween-20, and GluC protease was diluted in 100 mM ammonium bicarbonate. Proteolysis reactions were performed at a 1:250 Pif1:GluC (80ng GluC) ratio. To assess proteolysis, 6 µg of protein was mixed with SDS-PAGE loading dye and chilled on ice at the indicated time points. The stopped reactions were ultimately electrophoresed on 10% SDS-PAGE gels for 45 min at 150 V and stained with Coomassie Brilliant Blue staining dye.

## ACKNOWLEDGEMENTS

This work was funded by grants from the National Science Foundation (1929346) and Indiana CTSI Core Pilot Grant to L.B. and National Health Institutes (1R35GM133437) and American Cancer Society (RSG-16-180-01-DMC) to M.L.B. We would like to thank Dr. Song Tan for sharing the Piccolo NuA4 (Esa1) expression construct. Additionally, we would like to acknowledge Dr. Amber Mosley, IUSM Proteomics Core and the IU Bloomington Proteomics Core for help with the mass spectrometry analysis.

## SUPPLEMENTARY FIGURES

**Supplemental Figure 1:**
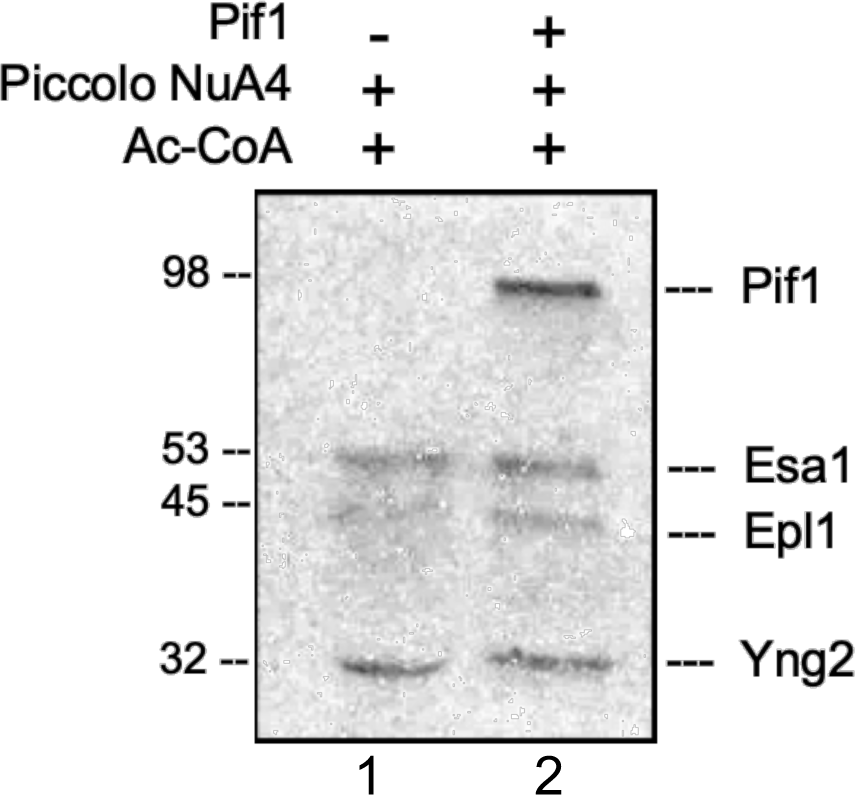
Piccolo NuA4 (Esa1) *in vitro-*acetylates Pif1: Recombinant NuA4 and full-length Pif1 were incubated along with ^14^C labelled-acetyl CoA as described in the Materials and Methods. The reaction products were separated on a 4-15% SDS-PAGE gel and subsequently subjected to autoradiography. Piccolo NuA4 (Esa1) was capable of robustly acetylating Pif1 *in vitro* (lane 2). The Esa1 subunit also underwent autoacetylation and acetylated the other subunits (Epl1 and Yng2) of the NuA4 complex (lanes 1 and 2).

**Supplemental Figure 2:**
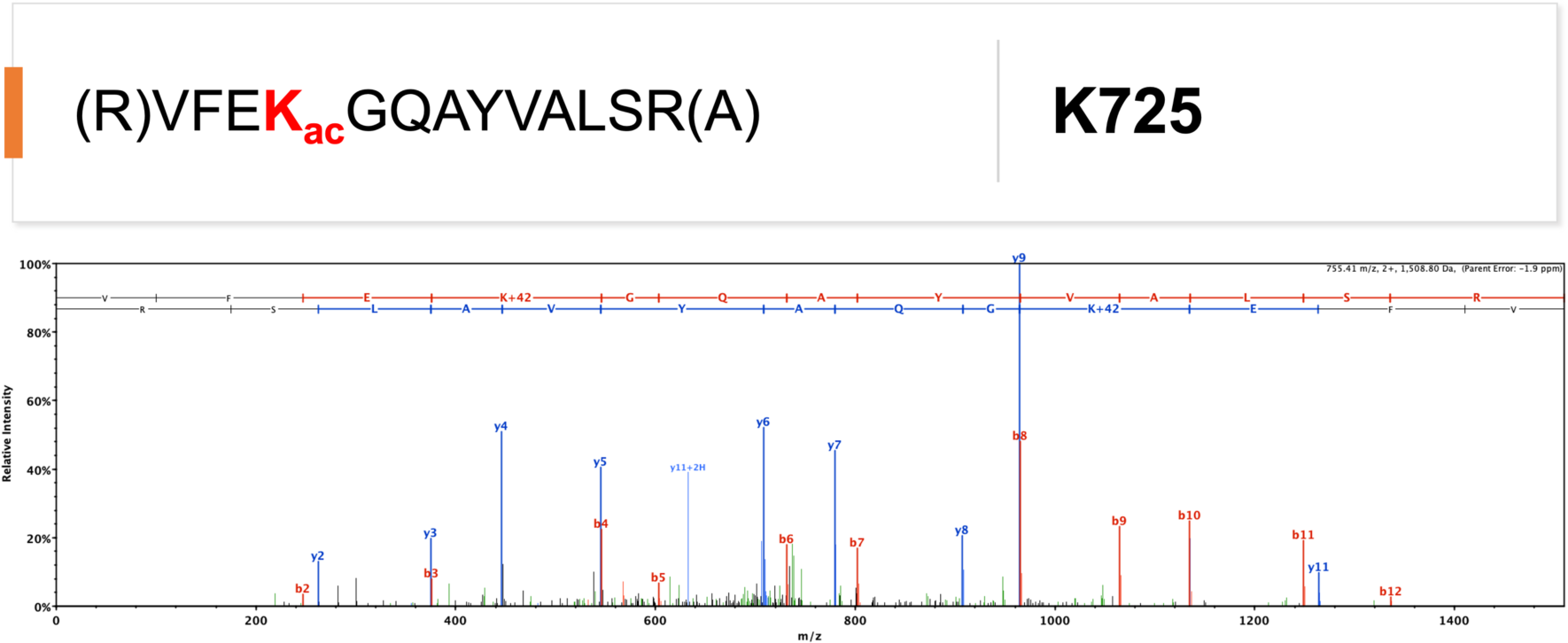
Spectra for Acetylated Pif1 Lysine 725: Representative spectra for lysine acetylation sites on Pif1 annotated on Scaffold (Proteome Software, Portland OR). The b-ions are labeled in red, and y-ions are labeled in blue. Neutral loss and other parent ion fragments are shown in green. The sequence of the acetylated peptide is denoted above the spectra with the acetylated lysine (K) highlighted in bold red font.

**Supplemental Figure 3:**
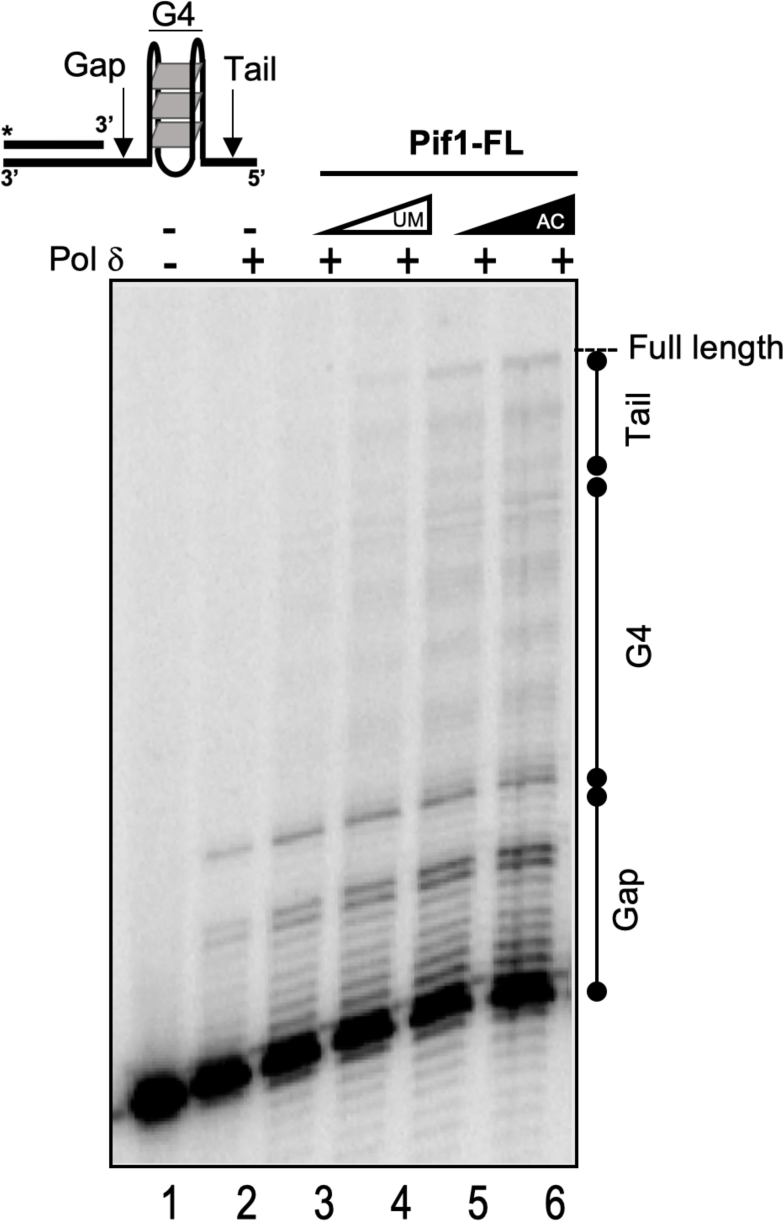
Pif1 resolves G4 structures to allow pol δ synthesis. The synthesis activity of 23 nM *S. cerevisiae* DNA polymerase delta (pol δ) was assayed on 5 nM cMyc-G4 substrate in the absence (lane 1) and presence of increasing concentrations (5 and 10 nM) of UM-Pif1 (lanes 3, 4) and AC-Pif1 (lanes 5, 6). The reactions were performed in a reaction buffer containing 20 mM Tris HCl (pH 7.8), 8 mM Mg(CH_3_COO)_2_, 100 mM KCl, 1 mM DTT, 0.1 mg/mL BSA, 100 μM dNTPs, and 1 mM ATP for 10 min at 30°C. Reactions were terminated using 2X termination dye and were immediately heated to 95°C and loaded onto a pre-warmed denaturing polyacrylamide gel (12% polyacrylamide, 7 M urea), and reaction products were separated by electrophoresis for 80 min at 80 W, subsequently dried, and analyzed. **Result**: DNA pol δ alone was unable to synthesize on the G4 substrate past the gap region, indicating the presence of a stable G4 structure. However, in the presence of both UM-Pif1 and AC-Pif1, we observed synthesis past the gap and into the G4 region. The AC-Pif1 displayed the highest stimulation of pol δ synthesis, including the formation of a full-length product, presumably because AC-Pif1 was more efficient at G4 structure resolution than UM-Pif1.

**Supplemental Figure 4:**
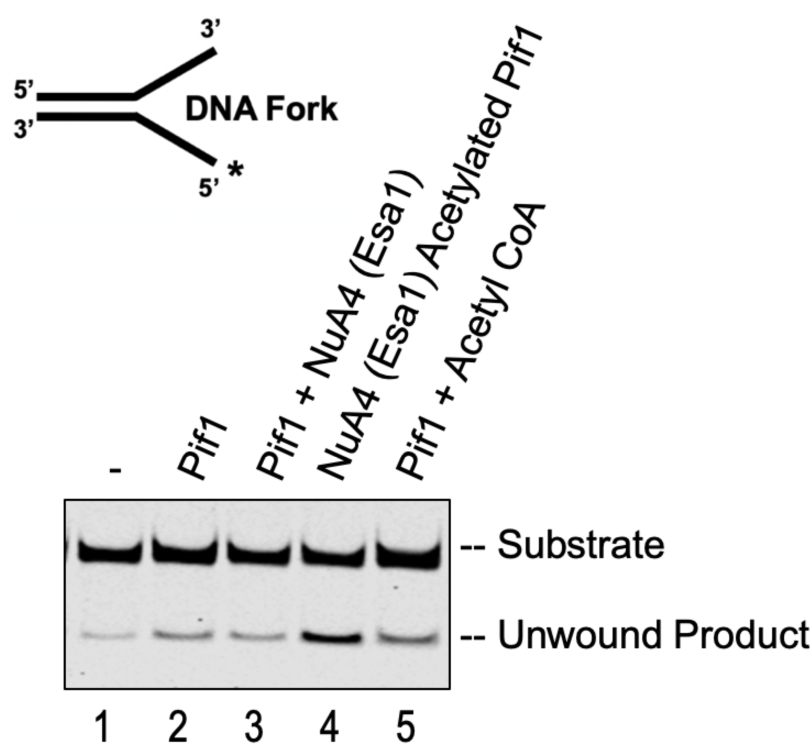
Stimulation of helicase activity is dependent on Pif1 acetylation alone. Helicase assays was performed using an IR labeled DNA fork in the presence of one nanomolar of either UM-Pif1 (lane 2), Pif1 + Piccolo NuA4(Esa1) (lane 3), AC-Pif1 (lane 4) or Pif1 + Acetyl CoA lithium salt as described in Materials and Methods.

## SUPPLEMENTARY TABLES

**Supplemental Table 1.**
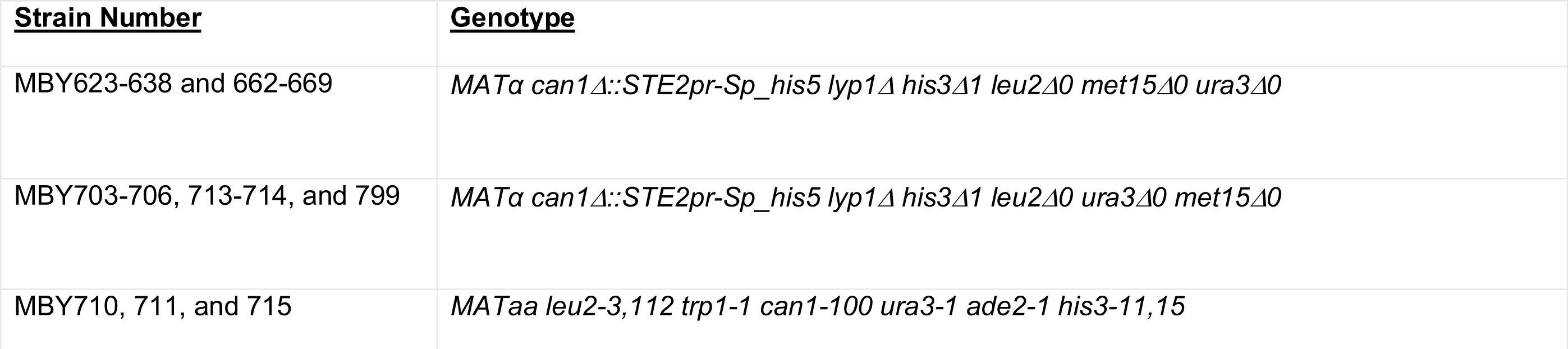
Yeast genotypes used in this study.

**Supplemental Table 2.**
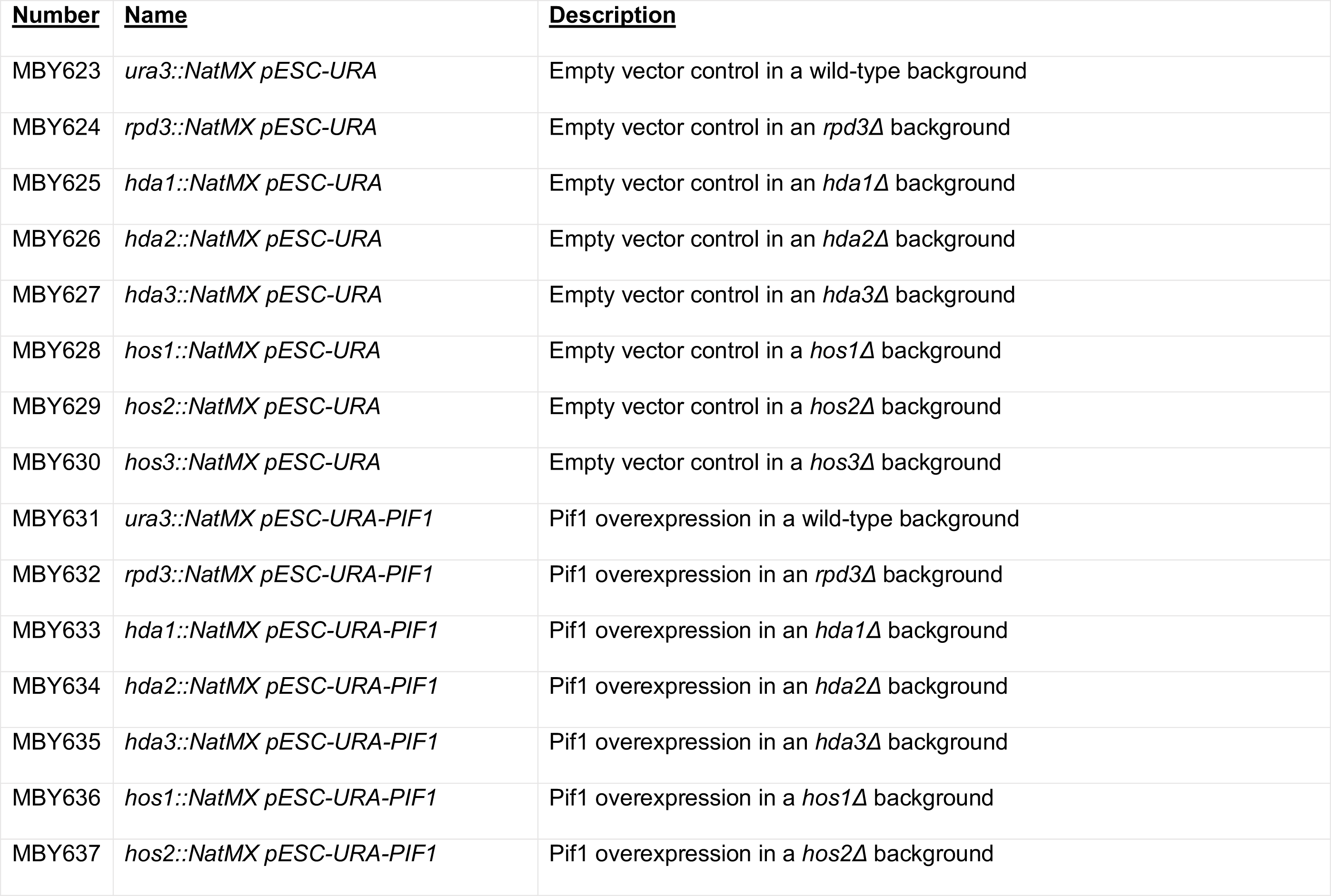
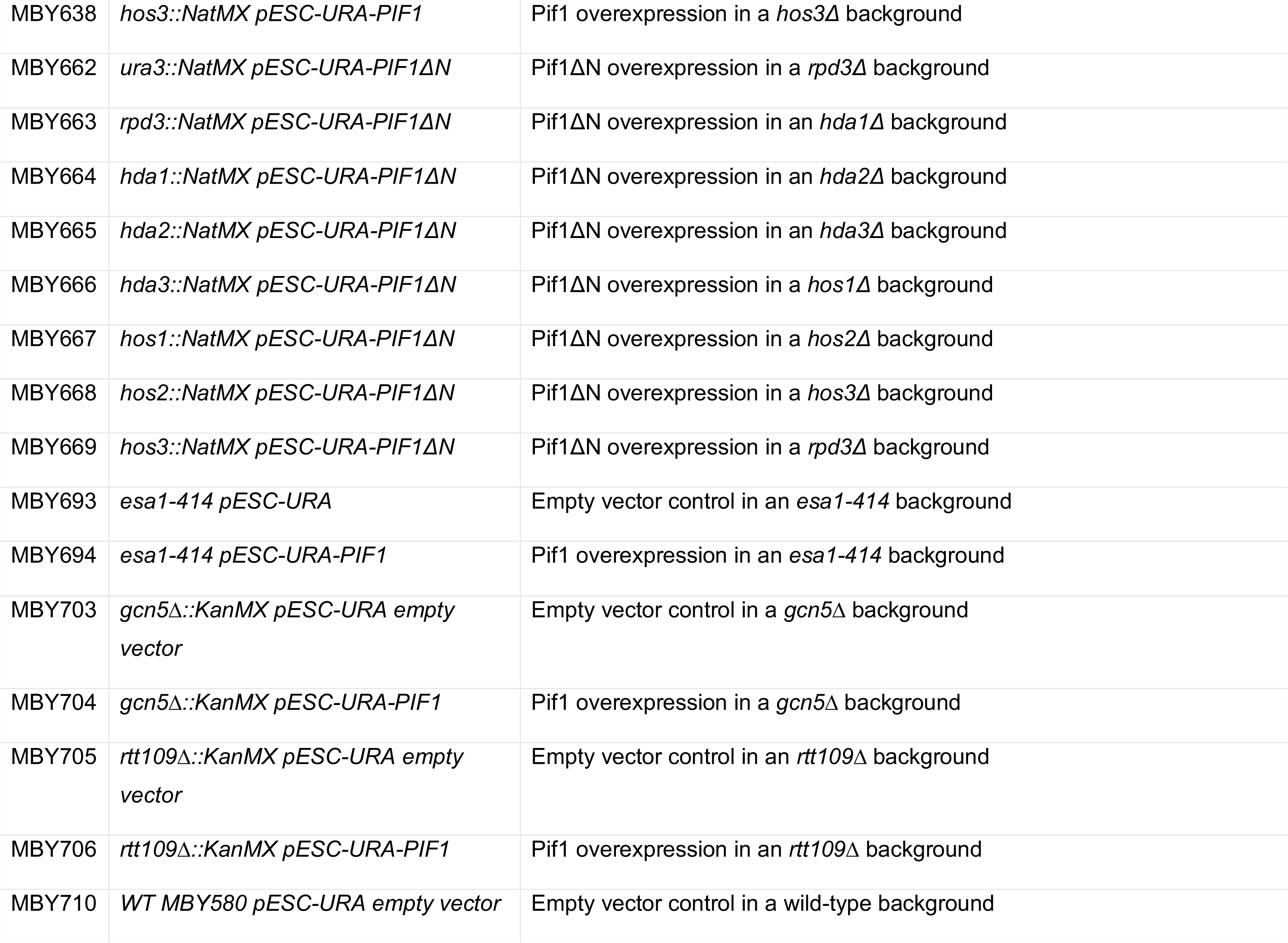
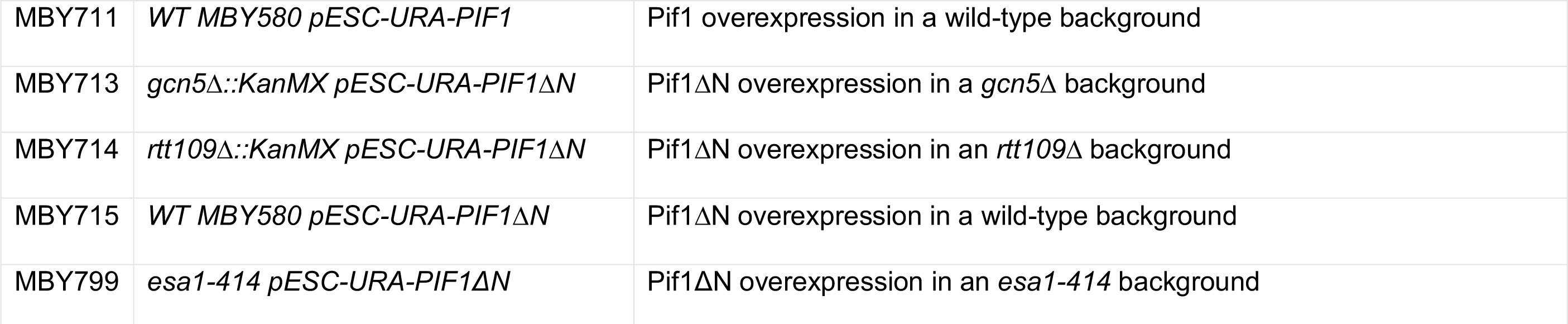
List of yeast strains used in this study.

**Supplemental Table 3.**
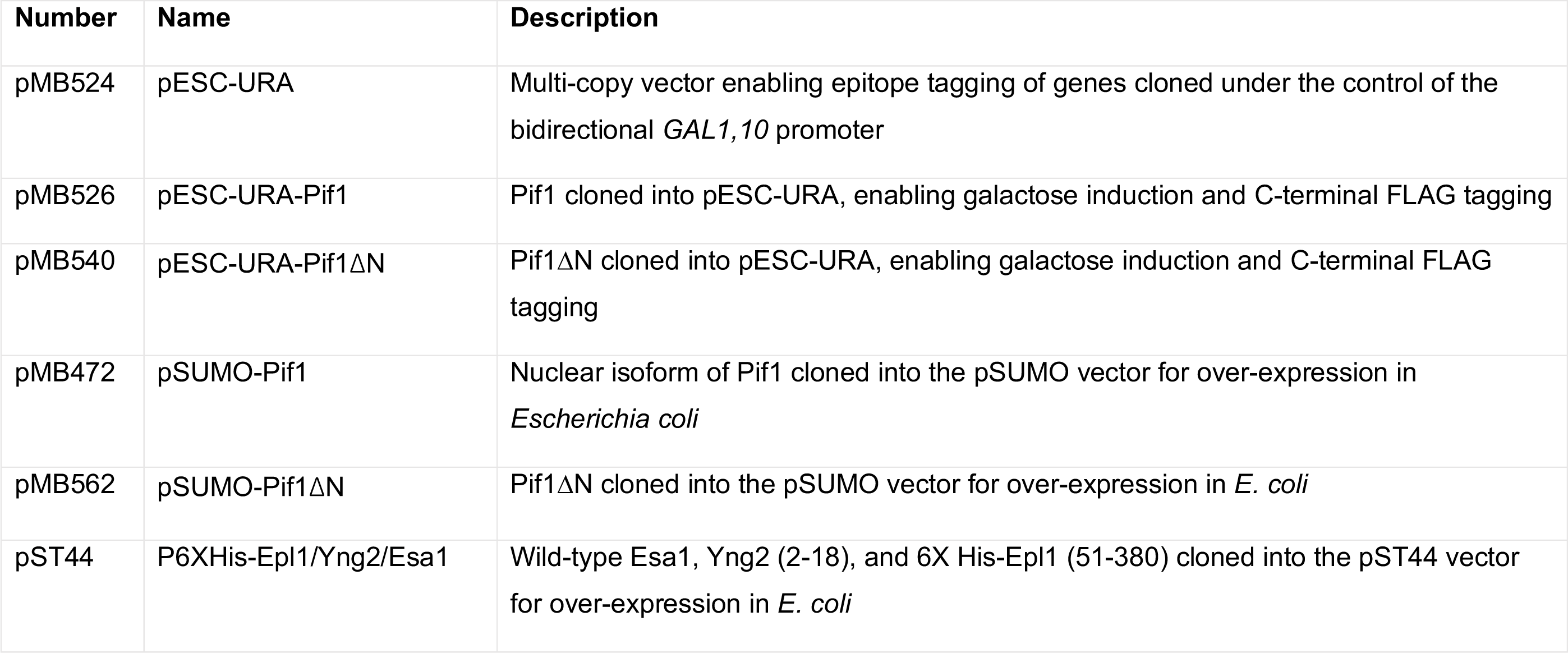
Plasmids used in this study.

| Number | Name | Description |
| --- | --- | --- |
| pMB524 | pESC-URA | Multi-copy vector enabling epitope tagging of genes cloned under the control of the bidirectional *GAL1,10* promoter |
| pMB526 | pESC-URA-Pif1 | Pif1 cloned into pESC-URA, enabling galactose induction and C-terminal FLAG tagging |
| pMB540 | pESC-URA-Pif1ΔN | Pif1ΔN cloned into pESC-URA, enabling galactose induction and C-terminal FLAG tagging |
| pMB472 | pSUMO-Pif1 | Nuclear isoform of Pif1 cloned into the pSUMO vector for over-expression in *Escherichia coli* |
| pMB562 | pSUMO-Pif1ΔN | Pif1ΔN cloned into the pSUMO vector for over-expression in *E. coli* |
| pST44 | P6XHis-Epl1/Yng2/Esa1 | Wild-type Esa1, Yng2 (2-18), and 6X His-Epl1 (51-380) cloned into the pST44 vector for over-expression in *E. coli* |

**Supplemental Table 4:**
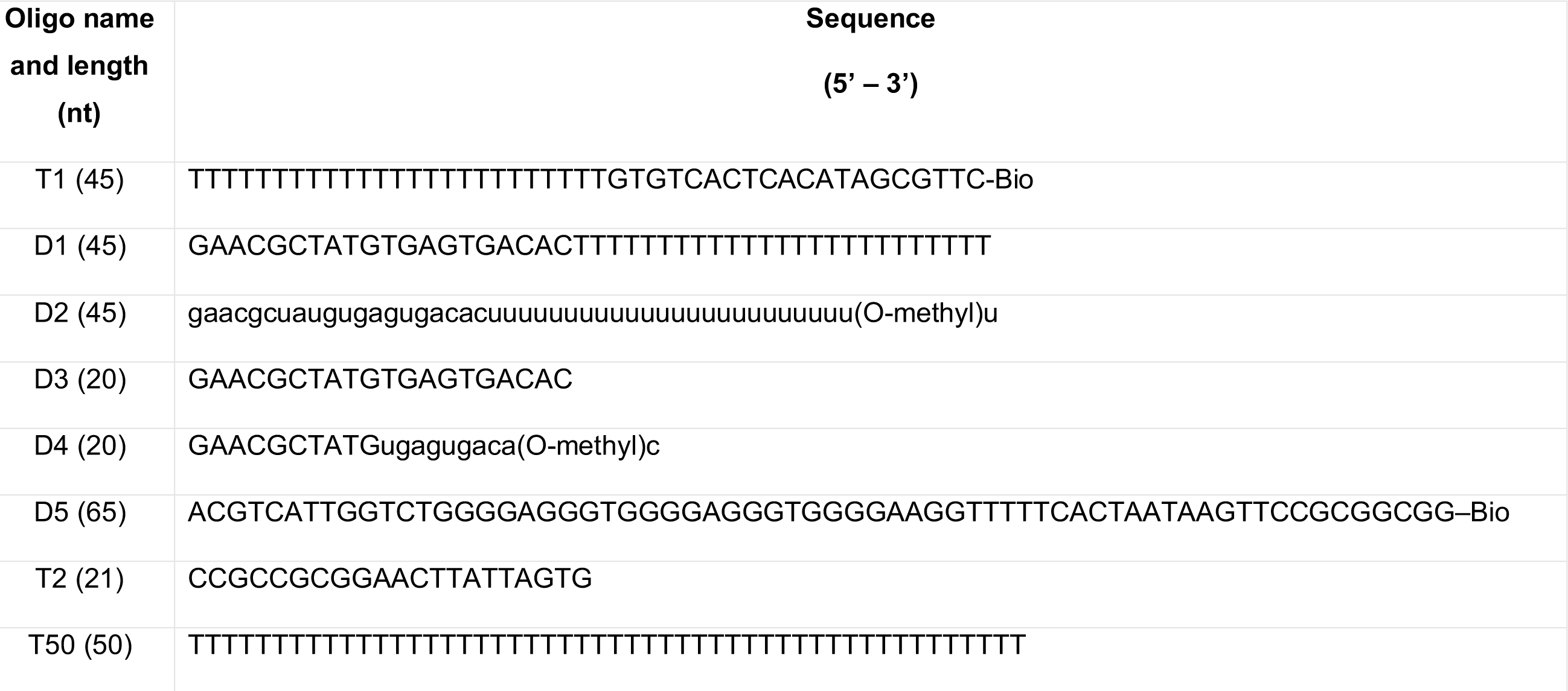
Oligonucleotides used in this study. Capital letters = dNTP, small letters = NTP, and Bio - biotinylated.

